# Targeting Ribosome Biogenesis as a Novel Therapeutic Approach to Overcome EMT-related Chemoresistance in Breast Cancer

**DOI:** 10.1101/2023.06.28.546927

**Authors:** Yi Ban, Yue Zou, Yingzhuo Liu, Sharrell B. Lee, Robert B. Bednarczyk, Jianting Sheng, Yuliang Cao, Stephen T. C. Wong, Dingcheng Gao

**Affiliations:** Department of Cardiothoracic Surgery, 1300 York Avenue, New York, New York 10065; Department of Cell and Developmental Biology, 1300 York Avenue, New York, New York 10065; Neuberger Berman Lung Cancer Center, 1300 York Avenue, New York, New York 10065; Sandra and Edward Meyer Cancer Center Weill Cornell Medicine, 1300 York Avenue, New York, New York 10065; Department of Radiology, Houston Methodist Hospital, 6565 Fannin Street, Houston, TX 77030; Department of Pathology and Laboratory Medicine, Houston Methodist Hospital, 6565 Fannin Street, Houston, TX 77030; Systems Medicine and Bioengineering Department Houston Methodist Cancer Center, Houston Methodist Hospital, 6565 Fannin Street, Houston, TX 77030

**Author notes:** These authors contributed equally (Yi Ban, Yue Zou).

**Keywords:** EMT, Epithelial-mesenchymal transition, EMT lineage tracing, ribosome biogenesis, RNA Polymerase I inhibitor, breast cancer, metastasis, chemoresistance

## Abstract

Epithelial-to-mesenchymal transition (EMT) contributes significantly to chemotherapy resistance and remains a critical challenge in treating advanced breast cancer. The complexity of EMT, involving redundant pro-EMT signaling pathways and its paradox reversal process, mesenchymal-to-epithelial transition (MET), has hindered the development of effective treatments. In this study, we utilized a Tri-PyMT EMT lineage-tracing model and single-cell RNA sequencing (scRNA-seq) to comprehensively analyze the EMT status of tumor cells. Our findings revealed elevated ribosome biogenesis (RiBi) during the transitioning phases of both EMT and MET processes. RiBi and its subsequent nascent protein synthesis mediated by ERK and mTOR signalings are essential for EMT/MET completion. Importantly, inhibiting excessive RiBi genetically or pharmacologically impaired the EMT/MET capability of tumor cells. Combining RiBi inhibition with chemotherapy drugs synergistically reduced metastatic outgrowth of epithelial and mesenchymal tumor cells under chemotherapies. Our study suggests that targeting the RiBi pathway presents a promising strategy for treating patients with advanced breast cancer.

**Significance:** This study uncovers the crucial involvement of ribosome biogenesis (RiBi) in the regulation of epithelial and mesenchymal state oscillations in breast cancer cells, which plays a major role in the development of chemoresistant metastasis. By proposing a novel therapeutic strategy targeting the RiBi pathway, the study offers significant potential to enhance treatment efficacy and outcomes for patients with advanced breast cancer. This approach could help overcome the limitations of current chemotherapy options and address the complex challenges posed by EMT-mediated chemoresistance.

## Introduction

Tumor cells exploit the transdifferentiation program of epithelial-to-mesenchymal transition (EMT) to acquire aggressive properties, including anchorage-independent survival, invasion, and stemness(1). Multiple growth factors (TGFβ, EGF, Wnts, etc), signaling pathways (Smad2/3, PI3K/Akt, ERK1/2, etc), EMT transcription factors (Snail, Twist, Zeb1/2, etc), and hundreds of downstream EMT related genes are involved in the EMT program(1). Such complexity leads to a wide spectrum of EMT phenotypes coexisting at different stages of tumors(2,3). The EMT-endowed features contribute to tumor heterogeneity, metastasis, and therapy resistance, making EMT an attractive therapeutic target.

Current EMT-targeting strategies focus on blocking EMT stimuli, signaling transduction, or mesenchymal features(2,3). However, these approaches may paradoxically promote the reversed process of EMT, mesenchymal to epithelial transition (MET), which also contributes to malignancy development(4,5). Therefore, we proposed that instead of targeting epithelial or mesenchymal phenotype, inhibiting a biological process mediating the transitions of both EMT and MET could effectively overcome the limitations of traditional strategies (2,3).

To investigate the EMT process in metastatic tumor progression, we previously developed an EMT lineage-tracing model (Tri-PyMT) by combining MMTV-PyMT, Fsp1(S100a4)-Cre, and Rosa26-mTmG transgenic mice (6). This model traces EMT via a permanent RFP-to-GFP fluorescence switch induced by mesenchymal-specific Cre expression. The absence of EMT reporting in metastatic lesions in this model has sparked a lively debate about the proper definition of EMT status in tumor cells and the biological significance of EMT in tumor progression (2,7). Rheenen’s group also posited that the Fsp1-Cre mediated EMT lineage tracing model might not accurately capture the majority of EMT events in comparison to an E cadherin-CFP model(8). Although we disagree with this assessment of Fsp1-Cre model’s fidelity in tracing EMT, we acknowledge the limitations of relying on a single EMT marker to investigate the EMT contributions in tumor metastasis. Notably, using the refined EMTracer animal models, Li et al. discovered that N-cadherin+ cells, rather than the Vimintin+ cells, were predominantly enriched in lung metastases(9). These findings with different mesenchymal-specific markers underscore the complex nature of the EMT process; metastases formation does not necessarily require the expression of many traditional mesenchymal markers.

The Tri-PyMT model, despite its limitations in comprehensive tracing of metastasis, provides unique opportunities to study EMT’s role in tumor progression and chemoresistance. In particular, the fluorescent marker switch of the established Tri-PyMT cell line reliably reports changes in EMT phenotypes(6,10). Using single-cell RNA sequencing (scRNA-seq) technology, we characterized the differential contributions of EMT tumor cells in tumor progression(10). Importantly, post-EMT(GFP+), mesenchymal tumor cells consistently demonstrated robust chemoresistant features compared to their parental epithelial cells (pre-EMT RFP+) (6), inspiring us to further study chemoresistance using the Tri-PyMT model.

## Materials and Methods

### Cell lines and cell culture

Tri-PyMT cells were established in our lab from primary tumors of MMTV-PyMT/Fsp1-Cre/Rosa26-mTmG transgenic mice(6). For experiments, we used Tri-PyMT cells from passage 5 (p5) to passage 10 (p10) which contains both RFP+ and GFP+ cells. MDA-MB231-LM2 cells were a gift from Dr. Joan Massague. To facilitate *in vivo* imaging, both cell lines were genetically labeled with luciferase by lenti-Puro-Luc. Cells were cultured in DMEM with 10% FBS, 1% L-Glutamine, and 1% Penicillin Streptomycin. A routine assay for Mycoplasma (Universal Mycoplasma Detection Kit, ATCC, Cat#30-1012K) was performed to avoid contaminations.

### Animals and tumor models

All animal works were performed in accordance with IACUC approved protocols at Weill Cornell Medicine. CB-17 SCID mice were obtained from Charles River (Wilmington, MA). For the experimental metastasis model, RFP+ and GFP+ Tri-PyMT cells were sorted from the 5-10^th^ passage cells and re-mixed at a ratio of 1:1. Total cells (1.5 ×10^5^ cells) were injected through the tail vein in 100 µL of PBS in 10-week-old females. For animals subjected to chemotherapy, Cyclophosphamide (CTX, 100mg/kg, Signa-aldrich, Cat# C0768) was administered once per week, i.p., for 3-4 weeks. BMH21 (25mg/kg, Sellechehem, Cat#S7718) was administrated 5 times/week, i.p., for 3-4 weeks. The progression of lung metastasis tumors was monitored by bioluminescent imaging (BLI) every 3-5 days on the Xenogen IVIS system coupled with analysis software (Living Image; Xenogen).

### Flow cytometry and cell sorting

For cultured cells, single-cell suspensions were prepared by trypsinization and neutralizing with the growth medium containing 10% FBS. For the metastatic lungs, cell suspensions were prepared by digesting tissues with an enzyme cocktail containing collagenase IV (1 mg/mL), hyaluronidase (50 units/mL), and DNase I (0.1 mg/mL) in Hank’s Balanced Salt Solution containing calcium (HBSS, Gibco) at 37°C for 20-30 minutes. Cells were filtered through a 40-μm cell strainer (BD Biosciences) and stained with anti-Epcam antibody (G8.8, Biolegend), if needed, following a standard immunostaining protocol. SYTOX Blue (Invitrogen) was added to the staining tube in the last 5 minutes to facilitate the elimination of dead cells.

Samples were analyzed using the BD LSRFortessa™ Flow Cytometer coupled with FlowJo_v10 software (FlowJo, LLC). GFP^+^ and RFP^+^ cells were detected by their endogenous fluorescence. Flow cytometry analysis was performed using a variety of controls including isotype antibodies, and unstained and single-color stained samples for determining appropriate gates, voltages, and compensations required in multivariate flow cytometry.

For fluorescence-activated cell sorting for *in vitro* culture or tail vein injection, we used the Aria II cell sorter coupled with FACS Diva software (BD Biosciences). The preparation of cells throughout sorting procedures was under sterile conditions. The purity of subpopulations after sorting was confirmed by analyzing post-sort samples in the sorter again.

### RNA-sequencing analysis

Total RNA was extracted from sorted RFP+, GFP+, and Double+ Tri-PyMT cells with the RNeasy Plus Kit (Qiagen). RNA-Seq libraries were constructed and sequenced following standard protocols (Illumina) at the Genomics and Epigenetics Core Facility of WCM. RNA-seq data were analyzed with customized Partek Flow software (Partek Inc). In brief, the RNA-seq data were aligned to the mouse transcriptome reference (mm10) by STAR after pre-alignment QA/QC control. Quantification of gene expression was performed with the annotation model (PartekE/M) and normalized to counts per million (CPM). Differential gene expression was performed with Gene Specific Analysis (GSA) algorithm, which applied multiple statistical models to each gene to account for its varying response to different experimental factors and different data distribution.

For heatmap visualizations, Z scores were calculated based on normalized per-gene counts. Algorisms for the biological interpretation of differential expressions between samples such as Gene Set Enrichment Analysis (GSEA) and Over-representation assay, are also integrated in Partek Flow platform. Gene sets of interests were downloaded from the Molecular Signatures Database (MSigDB, https://www.gsea-msigdb.org/gsea/msigdb.

### Single-cell RNA-sequencing (scRNA-seq) analysis

Single-cell suspensions were prepared following protocols from 10X Genomics. RFP+, GFP+, and Double^+^ Tri-PyMT cells were sorted by flow cytometry, and cells with >90% viability were submitted for sc-RNAseq at the Genomics and Epigenetics Core Facility at WCM. Single-cell libraries were generated using 10X Genomics Chromium Single-cell 3’ Library RNA-Seq Assays protocols targeting 8,000 cells from each fraction were sequenced on the NovaSeq sequencer (Illumina). The scRNA-seq data were analyzed with the Partek Flow software (Partek Inc), an integrated user-friendly platform for NGS analysis based on the Seurat R package(*11*). The raw sequencing data were aligned to the modified mouse transcriptome reference (mm10) containing RFP, GFP, PyMT, and Cre genes by STAR. The deduplication of UMIs, filtering of noise signals, and quantification of cell barcodes were performed to generate the single-cell count data. Single-cell QA/QC was then controlled by the total counts of UMIs, detected features, and the percentage of mitochondria genes according to each sample (**Supplementary Fig. S3A, S6A**). The top 20 principal components (PC) were used for tSNE visualization. Differential gene expression analysis, GSEA, Geneset Overrepresentation assay, and Cell Trajectory analysis (Monocle 2 model) are also integrated into the Partek Flow platform (Partek Inc).

To highlight the overall expression of feature genes in a pathway, such as ribosome biogenesis or EMT status, we calculated the AUCell values for the gene list. AUCell calculates a value for each cell by ranking all genes based on their expression in the cell and identifying the proportion of the gene list that falls within the top 5% of all genes. For the epithelial or mesenchymal gene lists, we selected genes based on their overall expression levels in Tri-PyMT cells, ensuring consistency with their reported associations to epithelial or mesenchymal phenotypes in the literature (**Supplementary Fig. S4A**). For RiBi genes, we used the ribosome biogenesis pathway gene list from the GOBP_Ribosome_Biogenesis (GO:0042254) in MSigDB.

### Tissue processing, Immunofluorescence, and Microscopy

Lungs with metastases were fixed in 4% paraformaldehyde overnight, followed by immersion in 30% sucrose for two days. They were then embedded in Tissue-Tek O.C.T. compound (Electron Microscopy Sciences). Serial sections (10-20 μm, at least 10 sections) were prepared for immunofluorescent staining.

For staining cultured or sorted cells, 2×10^3^ cells/well were seeded in 8-well chamber slides (Nunc Lab-Tek, Thermo Fisher) and cultured overnight in growth medium. Standard immunostaining protocols were followed using fluorescent-conjugated primary antibodies. Fluorescent images were captured using a Zeiss fluorescent microscope (Axio Observer) with Zen 3.0 software (Carl Zeiss Inc.).

### Western blot analysis

Cells were homogenized in 1x RIPA lysis buffer (Millipore) containing protease and phosphatase inhibitors (Roche Applied Science). The samples were then boiled in 1x Laemmli buffer with 10% β-mercaptoethanol and loaded onto 12% gradient Tris-Glycine gels (Bio-Rad). Western blotting was performed using the antibodies listed in the antibody table. Quantification of the Western blots was carried out using ImageJ. The relative intensity of each band was normalized to that of β-actin or tubulin, serving as loading controls for the same blot.

### Antibodies used in the experiments

**Antibodies used in the experiments:**
| Species | Antigen | Cat#, Company | Dilution | Application |
| --- | --- | --- | --- | --- |
| Mouse | E-Cadherin | 144725, Cell Signaling | 1:40 | WB |
| Mouse | EpCAM | G8.8 Biolegend | 1:100 | FC, WB,IF |
| Mouse | Vim | 5741T, Cell Signaling | 1:500 | WB |
| Mouse | β-actin | Ab8227, Abcam | 1:2000 | WB |
| Mouse | Fibrillalin | Ab5821, Abcam | 1:100 | IF |
| Mouse | p-ERK | 4370, Cell Signaling | 1:1000 | WB |
| Mouse | ERK | 4695, Cell Signaling | 1:1000 | WB |
| Mouse | p-mTORC1 | 3895s, Cell Signaling | 1:1000 | WB |
| Mouse | mTORC1 | 2983T, Cell Signaling | 1:1000 | WB |
| Mouse | p-rps6 | 4858T, Cell Signaling | 1:1000 | WB |
| Mouse | Tublin | 11224-1-AP, Proteintech | 1:1000 | WB |
| Mouse | Snail | 3895, Cell Signaling | 1:500 | WB |

### Cell viability assays

To determine the viability of Tri-PyMT cells under chemotherapy, cells (2×10^3^ cells/well) were seeded in 96-well adherent black-walled plates, and treated with a serial concentration of 5-FU, Cisplatin, 4-Hydroperoxy Cyclophosphamide, Doxorubicin, Gemcitabine, and Paclitaxel together with BMH21, for 72 hours. After treatment, cell viability was measured using the CellTiter-Glo® Luminescent Cell Viability Assay (Promega).

To quantify the cytotoxic effect and potential synergic effects of drug combinations, we used SynergyFinder 3.0(12) for data analysis. Basically, normalized cell viability data were formatted and analyzed with online server at SunergyFinder (https://synergyfinder.fimm.fi). The LL4 and ZIP methods were chosen for curve fitting and synergy calculation, respectively.

### Cell proliferation, transcription activity and nascent protein translation assays

To characterize cellular activities in nascent DNA, RNA, and protein synthesis, we applied EdU (5-ethynyl-2’-deoxyuridine), EU (5-ethynyl uridine), and OPP (O-propargyl-puromycin) incorporation assays, respectively. Cells (3×10^5^ cells/well) were seeded in a 6-well plate and treated (or left untreated) according to experimental settings. Cells were labeled with EdU (10 μM, for 30 min), EU (1 mM, for 1 hour), or OPP (10 μM, for 30 min) in the incubator. After labeling, cells were harvested by trypsinization, fixed with 3.7% formaldehyde in PBS, and permeabilized with 0.5% Triton-X100 in PBS. Labeling was detected using the Click-&-Go® detection kit (Vector Laboratories) following the standardized protocol and analyzed by flow cytometry.

### RT-PCR analysis

Total RNA was extracted from sorted RFP+, GFP+, and Doub+ Tri-PyMT cells using the RNeasy Plus Kit (Qiagen). For cDNA synthesis, 100 ng of RNA was used with the qScript cDNA SuperMix (Quanta Bio). Q-PCR was performed using SsoAdvanced™ Universal SYBR® Green Supermix (Bio-Rad) with target gene specific primers (as shown in the table) and Gapdh as the housekeeping control. The PCR protocol included an initial denaturation at 98°C for 2 minutes, followed by 40 cycles of 98°C for 15 seconds, 60°C for 30 seconds, and 72°C for 30 seconds, with signal readings at the end of each cycle. This was followed by a final extension at 72°C for 5 minutes and melt curve analysis on the CFX96 Real-Time System (Bio-Rad).

### Gene specific primers used in the experiments

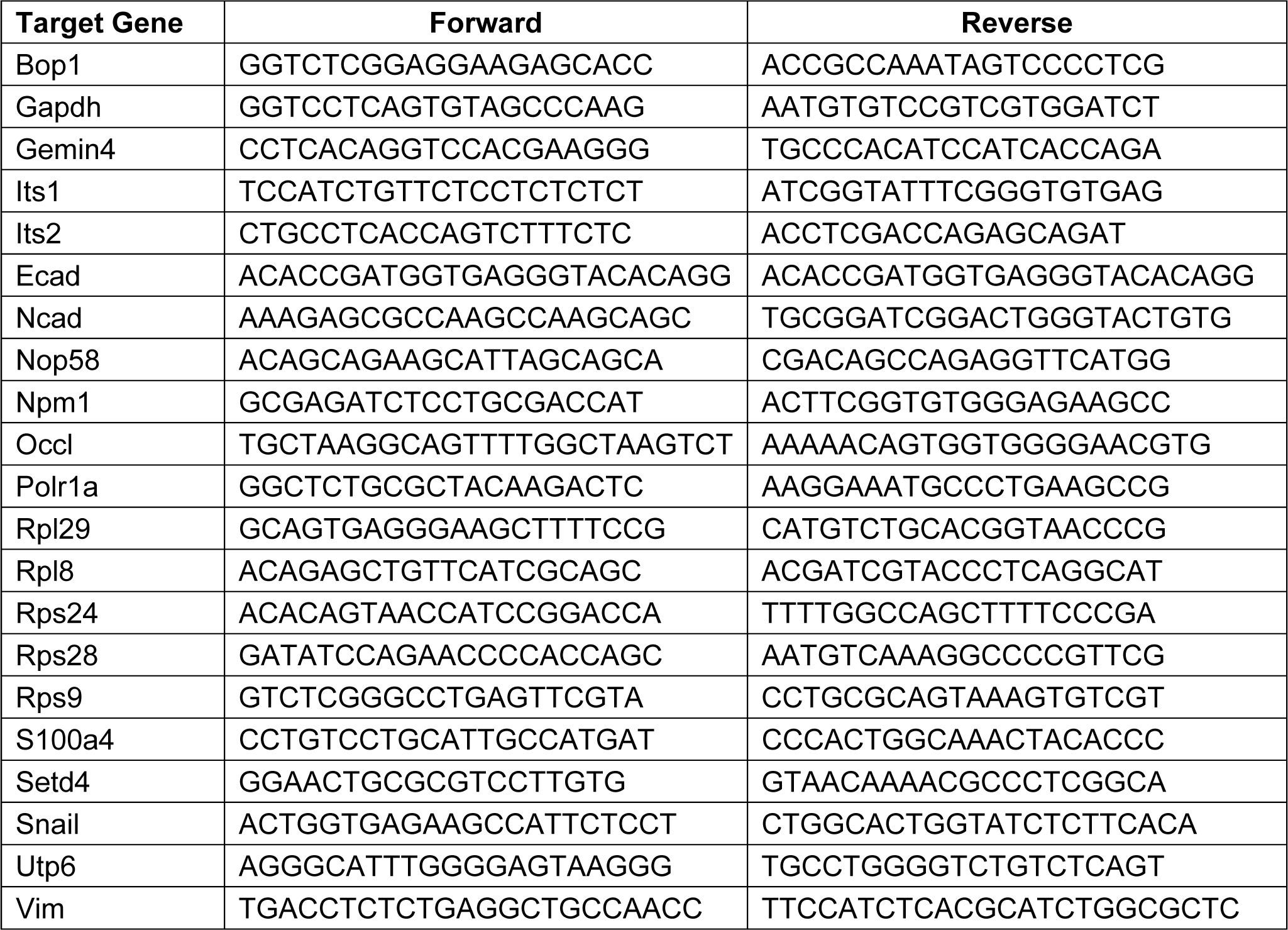
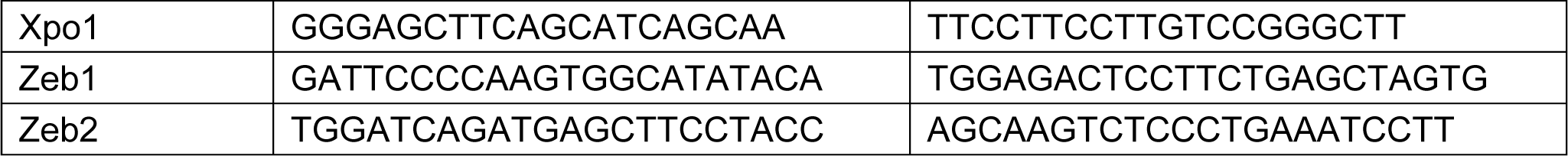

### Statistical Analysis

To determine the sample size for animal experiments, we performed power analysis assuming the (difference in means)/(standard deviation) is >2.5. Consequently, all animal experiments were conducted with ≥ 5 mice per group to ensure adequate power for two-sample t-test comparison or ANOVA. Animals were randomized within each experimental group, and no blinding was applied during the experiments. Results were expressed as mean ± SEM. Data distribution within groups and significance between different treatment groups were analyzed using GraphPad Prism software. P values < 0.05 were considered significant. Error bars represent SEM, unless otherwise indicated.

## Results

### Double+ Tri-PyMT cells mark the EMT transitioning phase

In contrast to their persistence in the epithelial state (RFP+) *in vivo* (6,10), RFP+ Tri-PyMT cells actively transition to GFP+ *in vitro* in growth medium containing 10% FBS (**Supplementary Fig. S1A**). As the fluorescent switch is irreversible, GFP+ cells accumulated over generations until a balanced ratio of RFP+/GFP+ is reached. This phenomenon indicates that the Tri-PyMT actively reports the ongoing EMT process in an EMT-promoting culture condition. Interestingly, we observed a subpopulation of cells, which were double positive for RFP and GFP, constituting approximately 2-5% of total cells (**Fig. 1A**). We posited that these RFP^+^/GFP^+^ (Doub^+^) cells represent tumor cells transitioning from an epithelial to a mesenchymal state, since their fluorescent marker cassette has switched to GFP expression induced by Fsp1-Cre, while pre-existing RFP protein lingers due to its tardy degradation. Indeed, immunoblotting analyses confirmed the association of the double positive fluorescence and a hybrid EMT status. Doub^+^ cells expressed intermediate levels of both epithelial markers (Epcam and E-cadherin) and the mesenchymal marker (Vimentin) (**Fig. 1B**). Further characterization of the Doub+ cells revealed higher percentages of S and G2/M phase cells in the Doub+ population compared to the RFP+ and GFP+ subpopulations (**Supplementary Fig. S1B**).

**Figure 1.**
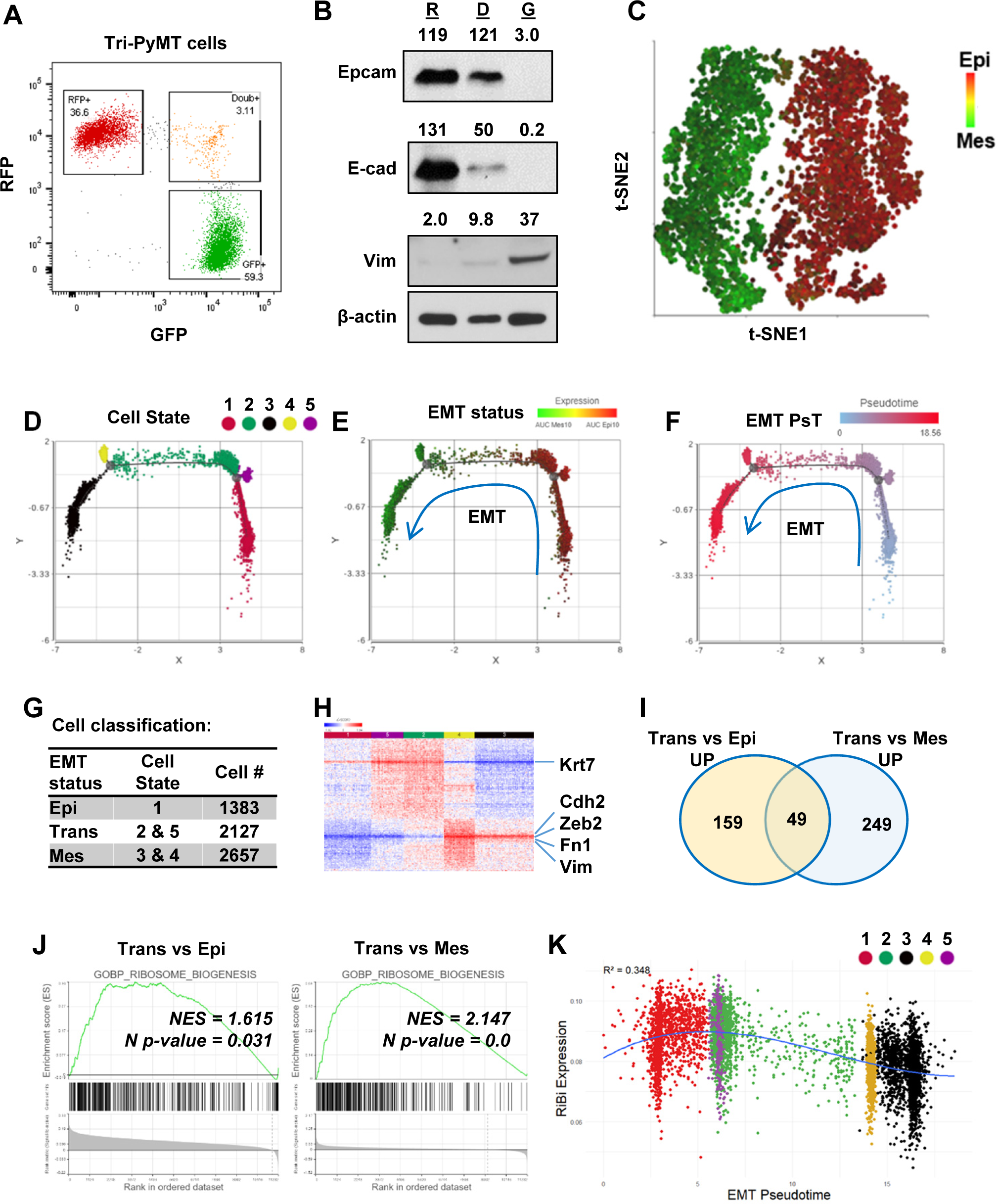
Activation of ribosome biogenesis pathway during the EMT transitioning phase in Tri-PyMT cells. **A**, Representative flow cytometry plot displays the percentage of RFP+, Double+ (Doub+, EMT transitioning cells), and GFP+ Tri-PyMT cells in culture. **B,** Western blot of EMT markers with flow sorted RFP+ (R), Doub+ (D), and GFP+ (G) Tri-PyMT cells. The Doub+ cells exhibit an intermediate EMT status with expression of epithelial marker (E-cadherin and Epcam) expression and higher mesenchymal marker (Vimentin) expression as compared with RFP+ cells. The numbers indicate normalized intensity of the band according to the β-actin band of the sample. **C,** The t-SNE plot of scRNA-seq analysis of Tri-PyMT cells. Two major cell clusters with differential EMT status are shown with the AUC value of 10 epithelial marker genes (in red) and 10 mesenchymal marker genes (in green). The marker genes are shown in Supplementary Fig S4A. **D, E, F,** Trajectory analysis using Monocle DDR tree. Cell trajectory analysis was performed with filtered EMT-related genes using Monocle 2 model. Five cell states (D) were identified with differential EMT statuses. From epithelial to mesenchymal phenotype, the Cell States were identified in order of 1, 5, 2, 4, and 3. EMT status (E) of cells was also highlighted by the AUC calculation with epithelial and mesenchymal marker genes. EMT Pseudotime (F) was calculated with Cell State 1 (the most epithelial state) as the root. **G,** Classification of cells with EMT states. Cells were classified according to their EMT state as Epi, Trans, and Mes; the cell number in each cluster is indicated in the table. **H,** The heatmap of differentially expressed genes in Trans cells compared to Epi and Mes cells. Totally, 313 genes were identified with criteria P <0.01, Fold change > 1.2, and Average expression >=5. **I,** The common enriched GO_BP pathways with GSEA when comparing Trans *vs.* Epi and Trans *vs.* Mes. There are 49 common pathways, including 5 pathways related to Ribosome Biogenesis (RiBi) or rRNA processing. Pathway names are shown in Supplementary Fig S4B. **J,** GSEA plots showing the specific enrichment of Ribosome Biogenesis (RiBi) pathway in EMT transitioning (Trans) cells compared to the Epi or Mes cells. **K,** The scatter plot displays the correlation of RiBi activity to EMT pseudotime. The polynomial regression line (order = 3) highlights the elevated RiBi pathway in Trans phase cells, R^2^ = 0.348, P < 0.001.

To further investigate the differential transcriptome in these EMT transitioning cells, we performed bulk RNA sequencing analysis using flow cytometry-sorted RFP^+^, Doub^+^, and GFP^+^ Tri-PyMT cells. Consistently, analysis of traditional EMT marker genes revealed that Doub+ cells expressed both epithelial and mesenchymal markers (**Supplementary Fig. S2A**). Differentially expressed genes in Doub+ cells were divided into 4 clusters, Trans_Up, Trans_Down, Epithelial and Mesenchymal markers (**Supplementary Fig. S2B**). Our attention was particularly attracted by upregulated genes within the Trans_Up cluster, as they may represent activated pathways specific in the transitioning phase of EMT. Interestingly, a gene set over-representative assay showed that the KEGG_Ribosome pathway was significantly enriched in the Trans_Up gene list (**Supplementary Fig. S2C**).

Together, these results indicate that Doub+ Tri-PyMT cells represent an active EMT-transitioning phase, as evidenced by well-established EMT markers; specific activations of biological processes in Doub+ cells warranrt further investigation for developing effective strategies to intervene the trasition.

### Ribosome biogenesis pathway is enhanced in the EMT transitioning phase

To gain a deeper understanding of transcriptome alterations and ensure adequate representation of EMT-transitioning cells, RFP^+^, GFP^+^, and Doub^+^ cells were sorted simultaneously via flow cytometry; equal numbers of each population were remixed and subject for single-cell RNA sequencing (scRNA-seq) analysis.

The *t*-SNE plot demonstrated two major clusters (**Fig. 1C):** one predominantly expressed epithelial genes, while the other displayed overall mesenchymal phenotypes. Doub+ cells were integrated into the two major clusters, suggesting that the overall single-cell transcriptome may not be sensitive enough to identify tumor cells at the EMT-transitioning phase. We therefore performed cell trajectory analysis (Monocle 2) based on all EMT-related genes (EMTome (13)). Five cell states related to EMT status were identified (**Fig. 1D**). All of them aligned well with a specific expression pattern of epithelial and mesenchymal marker genes based on AUC values (**Fig. 1E**), or individual epithelial/mesenchymal pairs, such as Fsp1/Epcam and Vim/Krt18 (**Supplementary Fig. S3B, S3C)**. Furthermore, we calculated the EMT pseudotime of each cell and designated State 1 (the most epithelial state) as the root (**Fig. 1F**). Cells were then classified into three main categories, Epi (State 1 cells), Trans (State 5&2) and Mes (State 4&3), based on their position within the EMT spectrum (**Fig. 1G**). Differential gene expression analysis confirmed that Trans cells gained expression of mesenchymal markers such as Cdh2, Vim, Fn1 and Zeb2, while retaining expression of epithelial markers such as Krt7 (**Fig. 1H**).

With the differential expression gene list, we analyzed the pathway activation in Trans cells by Gene Set Enrichment Analysis (GSEA) with the biology process (BP) gene sets of the GO term (14). Among the 49 overlapped gene sets that represented significantly upregulated pathways when comparing Trans vs. Epi and Trans vs. Mes (P <0.05), six pathways were related to the ribosome or rRNA processing (**Fig. 1I, Supplementary Fig. S4B**). These findings were in line with the previous bulk RNA sequencing analysis that indicated activation of RiBi pathway in EMT-transitioning (Doub+) cells. Consistently, GSEA using scRNAseq confirmed significant enrichment of RiBi genes in Trans cells compared to cells at Epi or Mes phase (**Fig. 1J**). We further mapped the EMT spectrum of cells based on their EMT pseudotimes or cell states, and correlated them with their RiBi activities. A significant trend of RiBi activation was observed in the trasitioning phase of EMT process (**Fig. 1K, Supplementary Fig. S3D**). It is worth noting that the RiBi activity is lowest in cells characterized with the latest EMT pseudotime, indicating that the elevation of RiBi activities is transcient during EMT (**Fig. 1K)**. RT-PCR analysis of also confirmed the relatively higher expression of RiBi related genes in Doub+ cells compared to RFP+ and GFP+ cells (**Supplementary Fig. S5**).

### Ribosome biogenesis pathway was upregulated during MET process

The transient elevation of RiBi during EMT prompted us to ask whether the MET process required the same. Since the fluorescence switch of Tri-PyMT cells is permanent, we tracked MET by monitoring the regain of epithelial marker by post-EMT (GFP+/Epcam-) cells. We sorted GFP^+^/Epcam^-^ Tri-PyMT cells via flowcytometry and injected them into *Scid* mice via tail vein. Approximately 50% of tumor cells expressed EpCam at 4 weeks post-injection, indicating active MET (**Fig. 2A**). We then sorted GFP^+^ tumor cells, including both EpCam^+^ and EpCam^-^ cells, from the lungs for scRNA-seq analyses.

**Figure 2.**
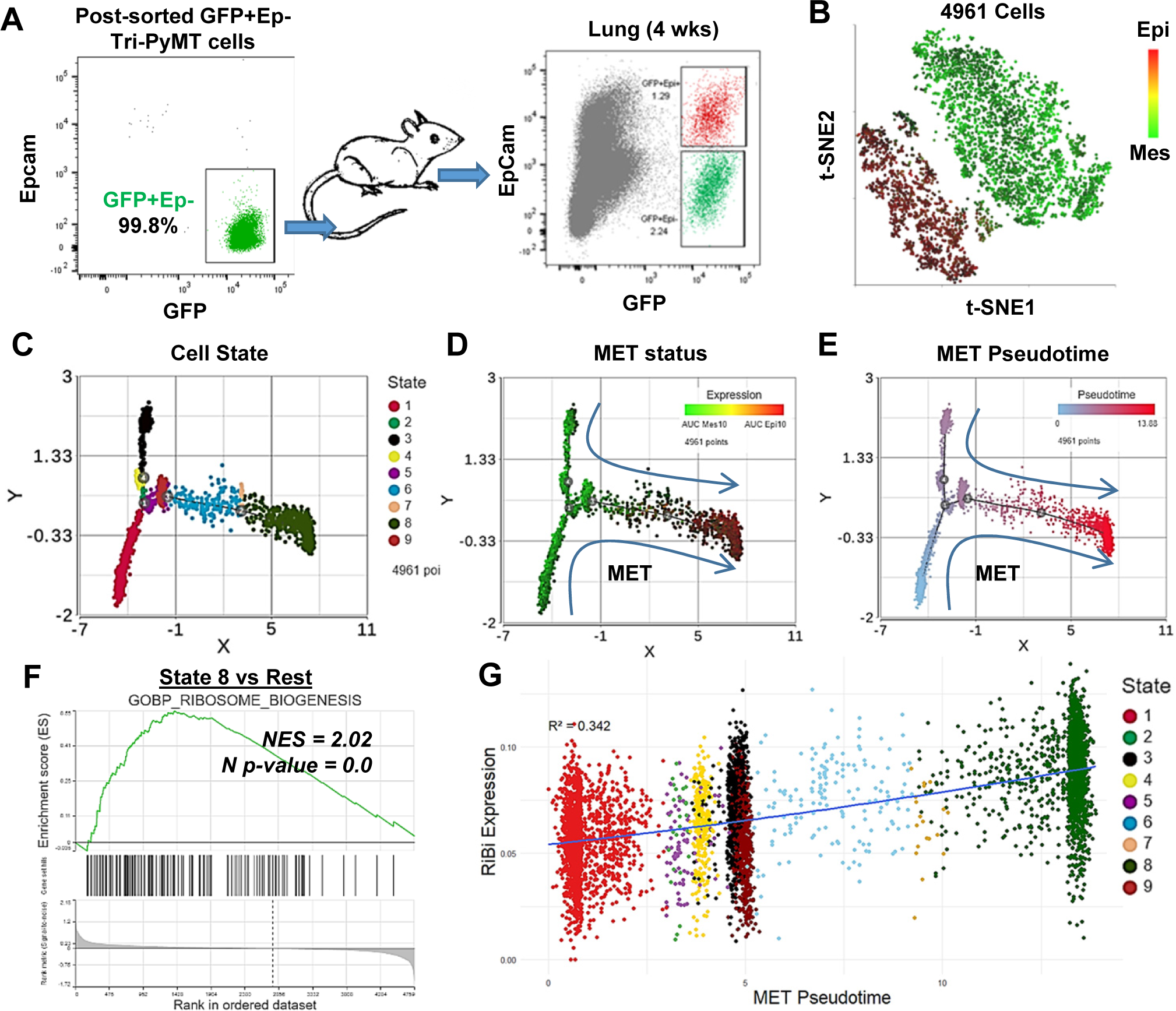
Activation of RiBi pathway in MET during lung metastasis outgrowth. **A**, Schematic of MET induction. GFP+ Tri-PyMT cells were sorted by flow cytometry (left) and injected into mice through the tail vein. Metastasis-bearing lungs were harvested after 4 weeks and analyzed for the gain of epithelial marker (EpCam) in tumor cells (right). Both Epcam+ and Epcam-cells were sorted and submitted for scRNA-seq analysis. **B,** The *t*-SNE plot of scRNA-seq analysis of GFP+ Tri-PyMT cells sorted from metastatic lungs of 3 individual animals. Two major cell clusters with distinct EMT statuses were identified, as indicated by the AUC values of 10 epithelial genes (in red) and 10 mesenchymal genes (in green). **C, D, E,** Monocle 2 model-based cell trajectory analysis identifies nine cell states (C). The MET process is emphasized by AUC values of mesenchymal and epithelial markers (D). MET Pseudotime is calculated with State 1 (the most mesenchymal state) as the root (E). **F,** GSEA plot exhibits the enrichment of the GOBP_Ribosome Biogenesis (RiBi) pathway in State 8 cells (the most epithelial phenotype) compared to the other states. **G,** Scatter plot illustrates the correlation between RiBi activity and MET pseudotime. A polynomial regression line (order = 3) emphasizes the elevated RiBi pathway during the MET process, R^2^ = 0.342, P < 0.01.

Similar to our observations with *in vitro* cultured Tri-PyMT cells, GFP+ cells from lungs formed two main clusters in the *t*-SNE plot, exhibiting overall epithelial or mesenchymal phenotypes (**Fig. 2B**). We performed trajectory analysis with EMTome genes and identified nine cell states with different EMT statuses (**Fig. 2C**). The relative expression of epithelial *versus* mesenchymal markers clearly illustrated the MET spectrum of tumor cells (**Fig. 2D, Supplementary Fig. S6B**). By designating the most extreme mesenchymal state (State 1) as the root, we calculated the MET pseudotime of individual cells (**Fig. 2E**). Consistent with the analysis for EMT, we found that tumor cells with epithelial phenotypes displayed elevated RiBi activity (**Fig. 2F, Supplementary Fig. S6C**), indicating its reactivation during MET process. A significant positive correlation was detected between RiBi gene upregulation and MET pseudotime (**Fig. 2G**). To eliminate the possibility that activation of RiBi pathway was solo related to the proliferation of cells, we project the S phase score to the scatter plot of Ribi activity/MET pseudotime. Indeed, cells in the far mesenchymal state show low S phase score, while the proliferating cells were mostly detected in the transitioning phase and epithelial phase (**Supplementary Figure S6D**). Together, these results suggested that upregulated RiBi pathway is equally needed for mesenchymal tumor cells to undergo MET during their outgrowth in the lungs.

### Activation of ERK and mTOR signaling pathways are linked to the upregulation of ribosome biogenesis

Ribosome biogenesis was recognized as a crutial factor in cancer pathogenesis a century ago (15). To further investigate the signaling pathways responsible for RiBi upregulation in the EMT/MET process, we sorted RFP^+^, Doub^+^ and GFP^+^ Tri-PyMT cells and probed the signaling activations in these cells. In response to serum stimulation, Doub^+^ cells exhibited significantly higher levels of phosphorylated extracellular signal-regulated kinase (p-ERK) and phosphorylated mammalian target of rapamycin complex 1 (p-mTORC1) compared to either RFP^+^ or GFP^+^ cells (**Fig. 3A**). The phosphorylation of Rps6, an essential ribosome protein of the 40S subunit, is regulated by synergistic crosstalk between mTORC1 and ERK signaling(16,17). We thus explored the status of p-Rps6 and observed a significantly higher level of p-Rps6 in Doub^+^ cells (**Fig. 3A**). These results imply a connection between differential signaling transductions and ribosome activities in EMT transitioning phase cells.

**Figure 3.**
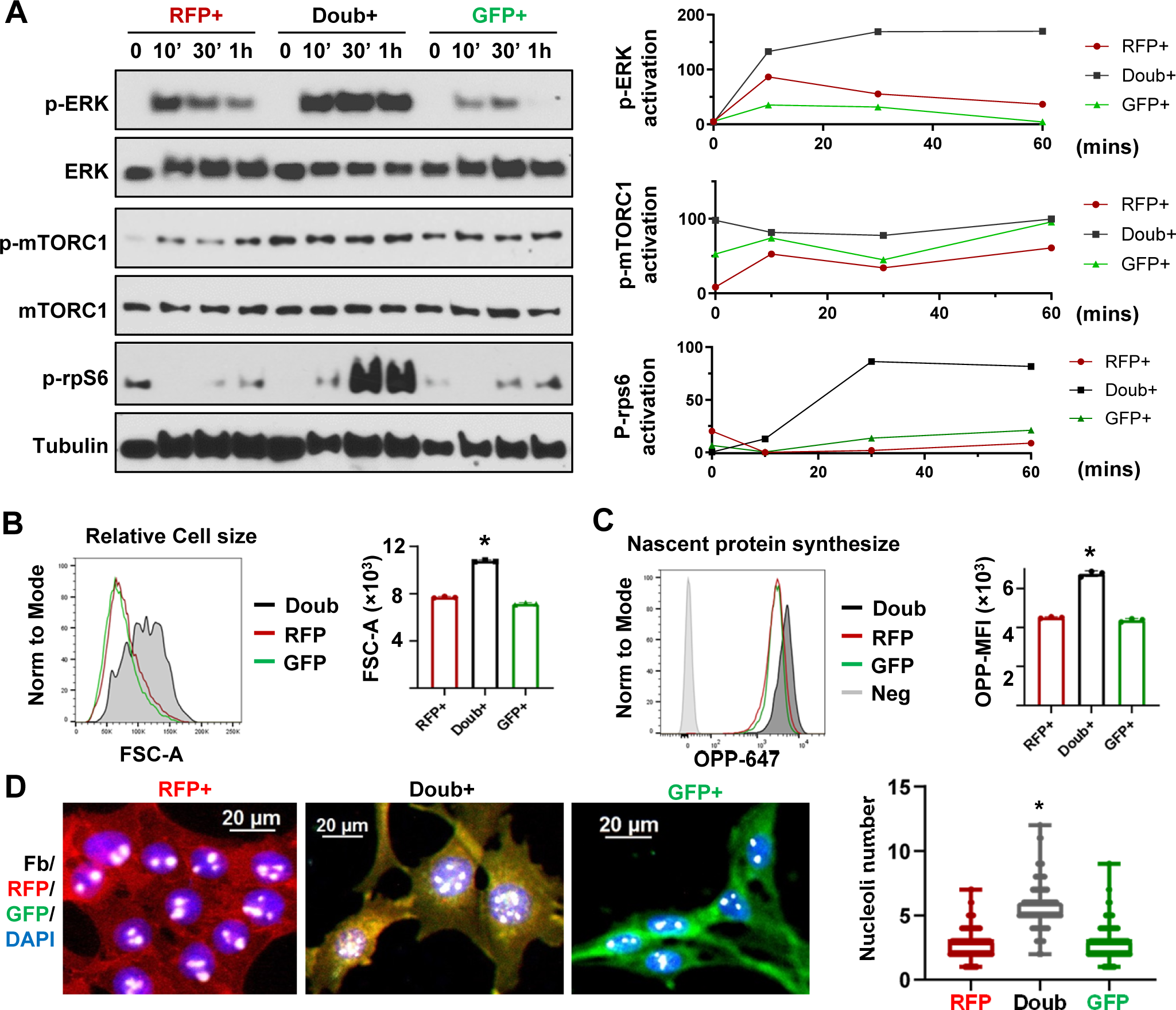
MAPK and mTOR pathways mediated ribosome biogenesis activation in EMT transitioning phase cells. **A,** Western blots show phospho-ERK, total ERK, phospho-mTORC1, total mTORC1 and p-rps6 in sorted RFP+, Doub+, and GFP+ Tri-PyMT cells following stimulation with 10% FBS at 0, 10, 30 and 60 mins. Activation of p-ERK, p-mTORC1 and p-Rps6 are quantified by the band intensity after normalizing to loading controls (right). **B,** Cell size analysis. Tri-PyMT cells (p5) were analyzed by flow cytometry. The relative cell size was indicated by FSC-A, for individual RFP+, GFP+, and Doub+ cells. Data from three biological replicates. **C,** Nascent protein synthesis assay. Histogram of OPP incorporation, as measured by flow cytometry, demonstrates enhanced nascent protein synthesis in Doub+ cells. The rate of new protein synthesis is assessed by adding fluorescently labeled O-propargyl-puromycin (OPP). Three biological replicates, One-way ANOVA, *P < 0.0001, Doub+ vs RFP+, and Doub+ vs GFP+. **D,** Fluorescent images reveal increased RiBi activity in Doub+ Tri-PyMT cells, as evidenced by Fibrillarin staining. RFP+, Doub+, and GFP+ Tri-PyMT cells were sorted and stained for nucleoli using an anti-Fibrillarin antibody. The quantification of nucleoli per cell is shown on the right. n = 522 (RFP+), 141 (Doub+), 289 (GFP+). One-way ANOVA, *P < 0.0001.

The primary function of ribosome is to synthesize proteins to support essential biological functions(18). Indeed, enhanced ribosome activity in Doub+ cells translated into distinct phenotypes in cell growth and nascent protein synthesis. Flow cytometry analysis showed that Doub+ cells possessed enlarged cell sizes compared to RFP^+^ or GFP^+^ cells (**Fig. 3B**). Using the O-propargyl-puromycin (OPP) incorporation assay, we found that Doub^+^ cells exhibited an increased rate in nascent protein synthesis (**Fig. 3C**). Another hallmark of ribosome activity is rRNA transcription that occurs in nucleoli. By immunostaining of Fibrillarin, a nucleolar marker(19), we found that the Doub^+^ cells had significantly more nucleoli compared to cells in epithelial (RFP^+^) and mesenchymal (GFP^+^) states (**Fig. 3D**). Consistently, using EU incorporation assay, we found significantly higher activity of transcription in Doub+ cells compared to RFP+ and GFP+ cells (**Supplementary Figure S7**).

Collectively, these results suggest that the elevation of RiBi pathway in cells at EMT transitioning phase is associated with aberrant activation of ERK and mTOR signalings, which may, in turn, confer nascent protein synthesis capability to tumor cells for completing phenotypic changes.

### Suppression of ribosome biogenesis reduced the EMT/MET capability of tumor cells

The transcription of ribosomal RNA (rRNA) is mediated by RNA polymerase I (Pol I) in eukaryotic cells (20). Small molecules such as BMH21 and CX5461 are specific Pol I inhibitors, which inhibit rRNA transcription and disrupt ribosome assembly(21,22), providing a specific RiBi targeting strategy. We then evaluated whether these Pol I inhibitors would affact EMT/MET process.

Fluorescence switch from RFP+ to GFP+ of TriPyMT cells were employed to investigate the impact of Pol I inhibitors on EMT. RFP+/Epcam+ cells were sorted via flowcytometry and served as cells in Epithelial state. In contrast to the vehicle-treated RFP^+^/Epcam+ cells, which transitioned to GFP^+^ upon serum stimulation, most cells treated with either BMH21 or CX5461 stayed in the RFP^+^ state (**Fig. 4A, Supplementary Fig. S8A**). Of note, the treatment of BMH21 or CX5461 indeed inhibited the transcription activity in cells as detected by EU incorporation assay (**Supplementary Fig 8B**). To confirm that the impaired fluorescence switch of RFP^+^ Tri-PyMT cells is associated with the retention of epithelial phenotypes, we performed immunoblotting of EMT markers. Indeed, these Pol I inhibitors significantly blocked the expression of mesenchymal markers, including Vimentin and Snail, while preserving the expression of the epithelial marker (Ecadherin) (**Fig. 4B**).

**Figure 4.**
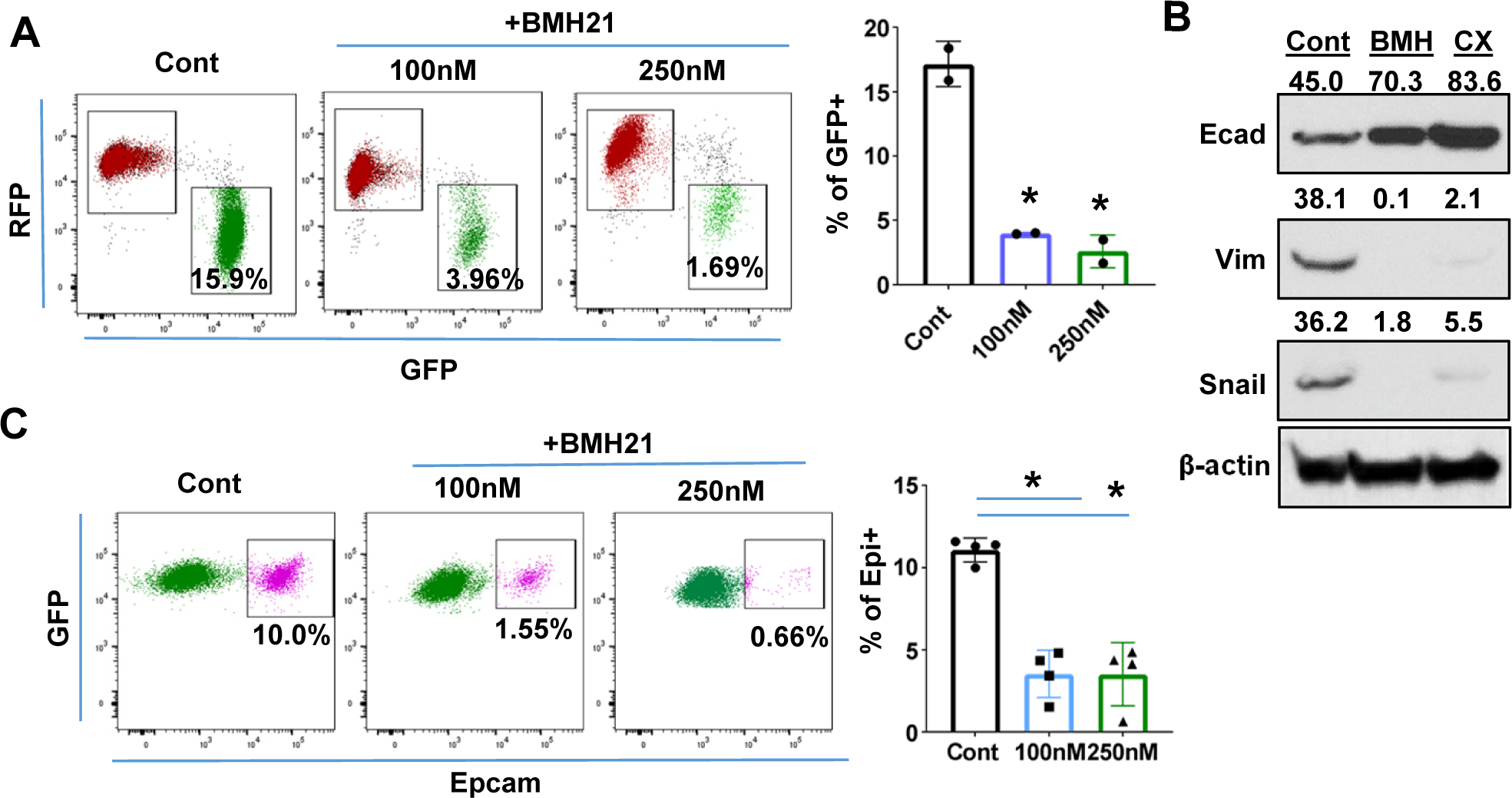
RiBi inhibition by Pol I inhibitor reduces the EMT/MET capability of tumor cells. **A,** Flow cytometry plots display the percentage of GFP+ cells following BMH21 treatment. RFP+/EpCam+ Tri-PyMT cells were sorted and cultured in growth medium with or without BMH21 for 5 days. Percentage of GFP+ cells was analyzed by flow cytometry. Biological repeat, n=2, One-way ANOVA, *P=0.0039, 100nM vs Cont; *P=0.0029, 200nM vs Cont. **B,** Western blots reveal the expression of the epithelial marker (E-cad) and mesenchymal markers (Vim and Snail) in Tri-PyMT cells following a 5-day treatment with BMH21 (100nM) and CX5461 (20nM). The numbers indicate normalized intensity of the band according to the β-actin band of the sample. **C, F**low cytometry plots show the percentage of Epcam+/GFP+ Tri-PyMT cells treated with BMH21. GFP+/EpCam-Tri-PyMT cells were sorted and culture in 3D to induce MET for 14 days. The gain of Epcam expression was analyzed by flow cytometry. n=4, One way ANOVA, *P<0.0001, 100nM vs Cont; *P<0.0001, 200nM vs Cont.

Given that the elevated RiBi activity occurs during MET process as well, we further assessed the impact of Pol I inhibitor on the retrieval of epithelial markers by mesenchymal tumor cells. The regain of EpCam expression by the sorted GFP+/EpCam-Tri-PyMT cells in a 3D culture was measured via flow cytometry. BMH21 significantly prevented the Epcam retrieval by GFP+/Epcam-Tri-PyMT cells, suggesting the impaired MET capability upon treatment (**Fig. 4C**). These results laid a foundation for pharmacological inhibition of RiBi pathway to block EMT/MET of tumor cells.

The requirement of RiBi activity during EMT/MET transitioning was also demonstrated by genetically modulating ribosome proteins. As the organelle for protein synthesis, ribosome comprises 4 ribosomal RNAs and approximately 80 structural ribosomal proteins(15). Depleting one r-protein usually causes decreases of other r-proteins in the same subunit, and ultimately compromises the overall ribosome assembly(23). We, therefore, employed Lenti-shRNAs targeting Rps24 and Rps28, two essential genes of 40S subunit, to genetically modulate the RiBi pathway in Tri-PyMT cells (**Supplementary Fig. S9A**). Effective knocking-down of Rps24 or Rps28 significantly reduced the number of nucleoli (**Supplementary Fig. S9B**). Accompanying with the downregulated RiBi activities were the impeded EMT (RFP-to-GFP switch) and the similarly reduced MET (regain of Epcam) (**Supplementary Fig. S9C, D**). These results suggested that the elevated RiBi activities are critical for tumor cells to maintain their abilities to shift between epithelial and mesenchymal states.

### RiBi inhibition synergizes with chemotherapy drugs

To assess the overall impact of Pol I inhibitor on both epithelial and mesenchymal tumor cells, we treated unsorted Tri-PyMT cells (containing both RFP+ and GFP+ populations) with BMH21 for 7 days. Less accumulation of GFP+ cells was observed with BMH21 treatment compared with untreated controls (**Fig. 5A**). In contrast, treatment with the chemotherapy drug, cyclophosphamide (CTX), resulted in more GFP+ cells (**Fig. 5A**), consistent with our previous findings that chemoresistant features against CTX were acquired through EMT (6).

**Figure 5.**
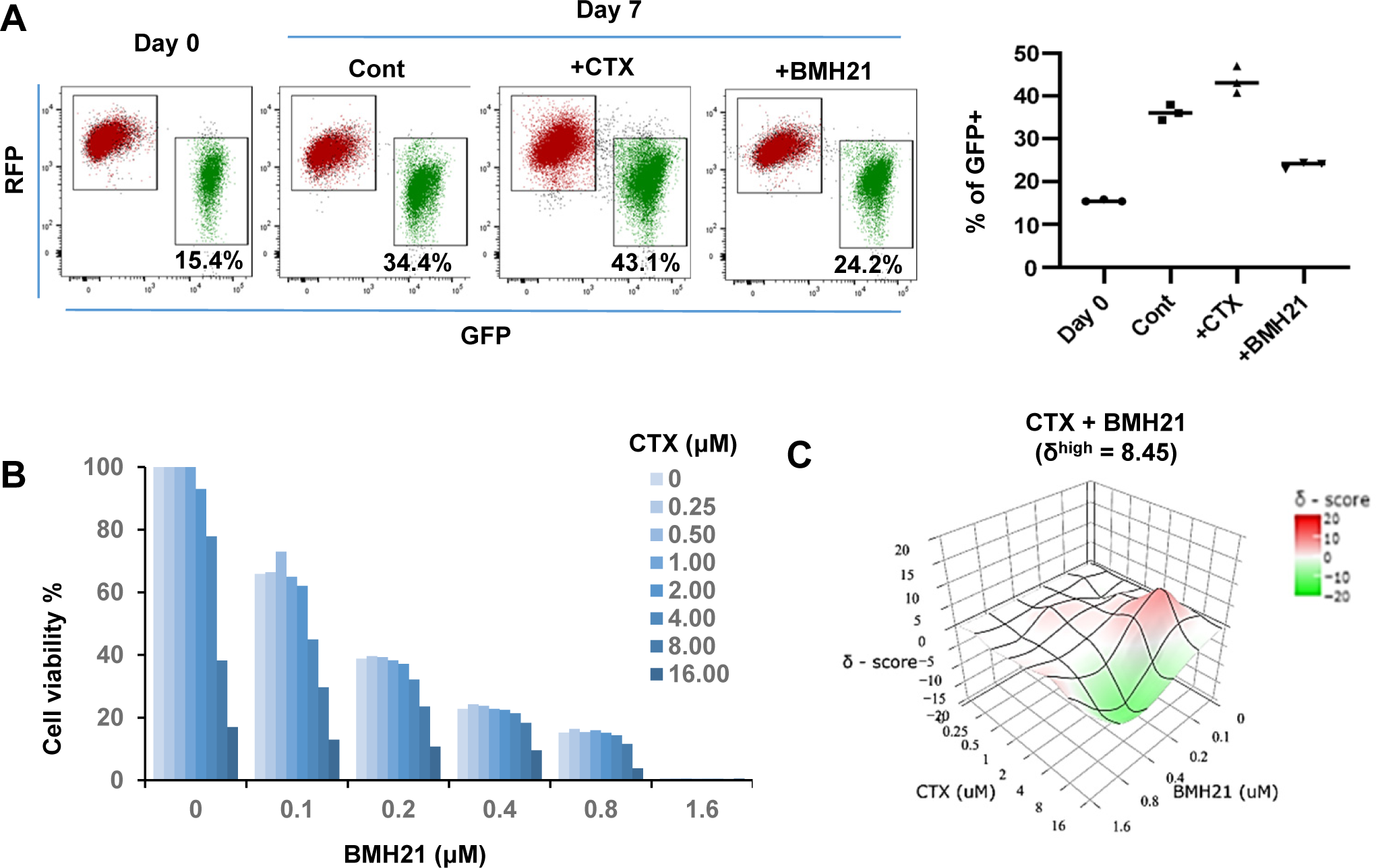
RNA Pol I inhibitor synergizes with chemo drug *in vitro*. **A,** Flow cytometry plots display the fluorescence switch of Tri-PyMT cells. Tri-PyMT cells (p5) were treat with cyclophosphamide (CTX, 2μM), Pol I inhibitor (BMH21, 0.1μM) or vehicle control for 7 days. Flow cytometry was performed to analyze the percentage of GFP+ cells. n=3 wells/treatment, One way ANOVA, *P=0.0050, Cont vs. CTX; *P=0.0002, Cont vs. BMH21. **B,** Cytotoxic assay of BMH21 and CTX treatments. Tri-PyMT cells were treated with serial concentrations of BMH21 (0, 0.1, 0.2, 0.4, 0.8, and 1.6 μM) in combination with serial concentrations of CTX (0, 0.25, 0.5, 1.0, 2.0, 4.0, 8.0, 16.0 μM). Bars represent the mean value of cell viabilities from duplicated treatments, n=2 wells/treatment. **C,** Synergy plots of BMH21 and Cyclophosphamide (CTX). Synergic scores (δ) for each combination were calculated using the ZIP reference model in SynergyFinder 3.0. Deviations between observed and expected responses indicate synergy (red) for positive values and antagonism (green) for negative values.

Based on these observations, we hypothesized that the blockade of EMT by Pol I inhibitor will reduce the EMT-mediated chemoresistance. The combination therapies of Pol I inhibitor and chemo drugs were therefore tested. We treated Tri-PyMT cells, which contain approximately 15% GFP+ cells, with a series of BMH21 concentrations with or without CTX. Cytotoxic assay after 3 days treatment revealed an enhanced sensitivity of tumor cells to the combination therapies (**Fig. 5B**). Interestingly, BMH21 and CTX exhibited optimal synergy (**Fig. 5C**). Particularly at concentrations of 100-200 nM, BMH21 showed significant high synergy scores with CTX (δ^high^ = 8.45). Such low concentrations of BMH21 were approximately 10-20% of the IC_max_ in Tri-PyMT cells and have also been shown to effectively block the EMT/MET process, suggesting the synergy was likely induced by EMT blockade, rather than cytotoxicity of BMH21. Importantly, the synergic effect between BMH21 and chemotherapy drugs is not limited to CTX. We performed the combination treatment of BMH21 with most commonly used chemodrugs of breast cancer therapy, including 5FU, Cisplatin, Doxorubicin, Gemcitabine and Paclitaxol. Trends toward synergies were also found with most of them, especially with lower concentrations of BMH21(100 – 200 nM) (**Supplementary Fig. S10**). These results suggest that RiBi blockade by the Pol I inhibitor may represent an effective approach to overcome EMT-related chemoresistance.

### RiBi inhibition diminished chemoresistant metastasis of breast tumor cells in the lung

Both the reduced EMT-mediated chemoresistance and the diminished MET during metastatic outgrowth upon BMH21 treatment encouraged us to evaluate the efficacy of combination therapy for treating animals bearing metastatic breast tumors.

We established a competitive metastasis assay by injecting an equivalent number of GFP^+^ and RFP^+^ Tri-PyMT cells (GFP:RFP, 1:1, representing epithelial and mesenchymal tumor cells, respectively) into *Scid* mice via the tail vein. Tumor-bearing mice received the vehicle, single, or combination therapy of BMH21 (25mg/kg, 5 times/week for 3 weeks, *ip.*) and CTX (100mg/kg, once a week for 3 weeks, *i.p.*). Bioluminescent imaging (BLI) revealed that the combination treatment exhibited the highest restraint on the chemoresistant outgrowth of metastatic tumors compared to the vehicle, BMH21, or CTX mono-treatment groups (**Fig. 6A**). To analyze the differential impacts on epithelial and mesenchymal tumor cells by the therapies, we quantified the residual RFP+ and GFP+ cells by flow cytometry analysis. Significant decrease of both RFP+ and GFP+ cells were observed in mice receiving the combination therapy of BMH21 and CTX (**Supplementary Fig. S11).** Microscopic analyses also revealed that both RFP^+^ and GFP^+^ cells grew into macrometastases in the lung of vehicle-treated animals (**Fig. 6B**). Notably, many GFP^+^ cells displayed epithelial markers, such as EpCam (**Fig. 6C**), indicating that a MET process was involved during the outgrowth of lung metastasis. CTX treatment eliminated the majority of RFP^+^/epithelial metastases, while the GFP^+^/mesenchymal cells showed survival advantages under chemotherapy, resulting in a higher ratio of GFP:RFP. BMH21 treatment inhibited metastatic tumor growth. Interestingly, most tumor cells under BMH21 treatment kept their EMT phenotypes as RFP^+^/Epcam^+^ or GFP^+^/Epcam^-^ (**Fig. 6C**), indicating the impaired EMT/MET transitioning by RiBi inhibition. Importantly, the combination of BMH21 and CTX eliminated most tumor cells including both RFP+ and GFP+ cells, and significantly inhibited the outgrowth of metastatic nodules under chemotherapy (**Fig. 6B-6C, and Supplementary Fig. S11**).

**Figure 6.**
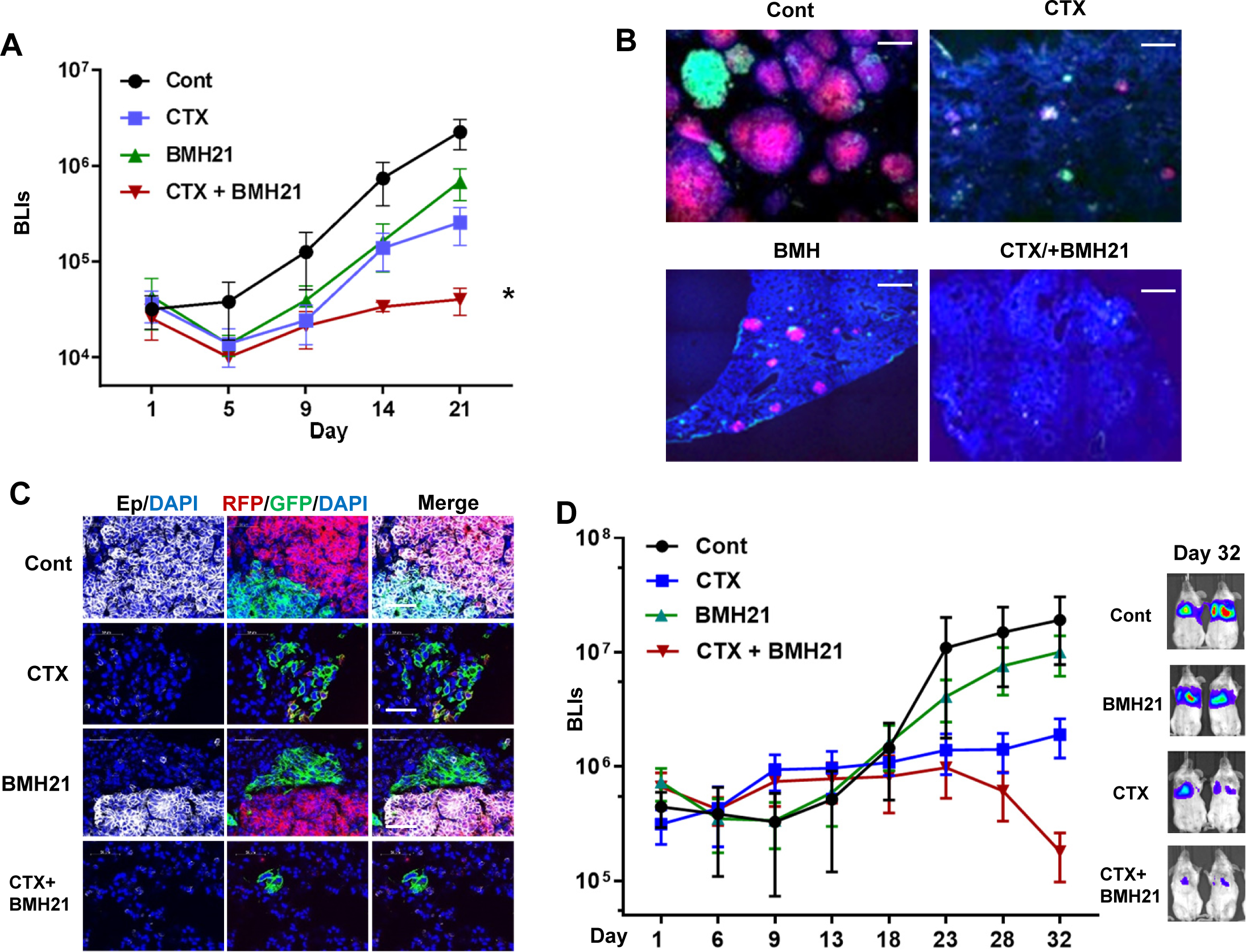
RNA Pol I inhibition synergizes with chemo drug in treating lung metastatic tumors. **A-C,** The same number of RFP+ and GFP+ Tri-PyMT cells were injected in animals via tail vein. Animals were treated CTX (100mg/kg, once a week, i.p.), BMH21 (20mg/kg, 5 times a week, i.p.) or both in combination. **A,** Lung metastasis growth curves as measured by bioluminescent imaging (BLI). n=5, two-way ANOVA with Tukey’s test, \**P*=0.0154, Cont *vs*. CTX+BMH; \**P*=0.0151, BMH21 *vs.* CTX+BMH21; \**P*=0.0372, CTX *vs.* CTX+BMH21 at Day 21 post-inoculation. **B,** Fluorescent images display RFP+ and GFP+ metastatic nodules in the lungs treated with CTX and BMH21. Cell nuclei were stained with DAPI. Scale bar: 200μm. **C,** Fluorescent images exhibit EMT statuses of RFP+ and GFP+ metastases. Sections were stained with anti-EpCam antibody; cell nuclei were stained with DAPI. Scale bar: 50μm. **D,** Lung metastasis growth curves for LM2 model. Lung metastatic breast tumor cells (LM2 cells) were injected into animals via tail vein. Animals were treated CTX (100mg/kg, once a week, i.p.), BMH21 (20mg/kg, 5 times a week, i.p.) or both in combination. n=5, two-way ANOVA with Tukey’s test, \**P*=0.0404, Cont *vs.* CTX+BMH21; \**P*=0.0161, BMH21 *vs.* CTX+BMH21; \**P*=0.0187, CTX *vs.* CTX+BMH at Day 32 post-inoculation. Representative BLI images of Day 32 were presented on the right.

We further established an experimental lung metastasis model with human breast cancer cells (MDA-MB231-LM2). The LM2 cells predominantly exhibited a mesenchymal phenotype *in vitro* and gained Ecad expression during the outgrowth of lung nodules *in vivo* (**Supplementary Fig. S12)**, indicating the involvement of the MET process. Consistent with the Tri-PyMT model, BMH21 synergized with CTX, leading to a significantly lower metastatic LM2 tumor burden than the mock or mono-treatment groups (**Fig. 6D**).

To further investigate the potential association of RiBi activity with EMT status of tumor cells in human breast cancer, we analyzed scRNA-seq data of primary tumor cells from two breast cancer patients (GSE 198745). A trend of relatively higher RiBi activity was detected in tumor cells with lower EMT pseudotime (**Supplementary Fig 13**). Interestingly, a pattern showing the highest RiBi activity in EMT/MET transitioning phase was detected in the sample of patient B (**Supplementary Fig 13F**), indicating the similar role of RiBi upregulation in EMT status of breast cancer patients. To examine whether RiBi activity associated with clinical outcomes of breast cancer patients, we analyze the RNAseq data in cBioPortal databases (www.cbioportal.org), including the TCGA PanCancer Atlas (1084 samples) and METABRIC (2500 samples). RiBi activities of tumors were quantified by the average z-scores of genes in RiBi pathway (303 genes). Patient samples with a score >1 were denoted as RiBi^High^, while patients with a score <0.5 were denoted as RiBi^Low^. The analyses of the survival data showed a significantly worse prognosis in the RiBi^High^ group compared to the RiBi^Low^ group (**Supplementary Fig 14**).

In summary, these results suggest that inhibition of RiBi activities diminished EMT/MET transitioning capability of tumor cells. In combination with chemotherapy drug, RiBi inhibition significantly reduced the outgrowth of chemoresistant metastasis. These results suggested that targeting RiBi-mediated EMT/MET process may provide a more effective therapeutic strategy for advanced breast cancer.

## Discussion

Targeting EMT for cancer therapy has been challenging due to the complexity of the EMT process and controversies of promoting MET which also favors tumor progression. Using the EMT-lineage-tracing model, we found an upregulation of RiBi pathway at the transitioning phase of both EMT and MET (**Supplementary Fig 15**). The transcient activation of RiBi pathway during the EMT process has been reported previously. Prakash et al. found that elevated rRNA synthesis/RiBi pathway was concomitant with cell cycle arrest induced by TGFβ, fueling the EMT program in breast tumor cells(24). Using the Tri-PyMT model, we found that the EMT transitioning (Doub+) cells had higher percentages of cells in the S and G2/M phases compared to the RFP+ and GFP+ cells. This is inconsistent with the observation that RiBi activity was higher in G1 arrested cells treated with TGFβ (24). This discrepancy may be due to the different EMT stimuli used in the experiment systems. Additionally, further investigations are needed to determine whether a cell could complete the EMT process within a single cell cycle or requires multiple cell divisions.

A more recent study identified a subpopulation of circulating tumor cells (CTCs) in which high RiBi activities persisted to maintain their high metastatic potentials(25). Interestingly, the RiBi activity was associated with epithelial phenotypes rather than mesenchymal ones in CTCs(25). Indeed, EMT induction by TGFβ primarily suppressed ribosome gene expression and global translational activity (25). By employing our unique EMT-lineage-tracing model, we discovered that the RiBi pathway was transiently elevated during the transitioning phases of EMT/MET program. The enhanced activation of RiBi pathway diminished as tumor cells accomplished phenotype changes. In general, a lower RiBi activity was observed in the mesenchymal tumor cells as compared to that in the epithelial ones. Importantly, the involvement of unwonted RiBi activities during both EMT and MET processes makes RiBi a new and better target for overcoming EMT-related chemoresistance and chemoresistant metastasis.

Targeting RiBi pathway by RNase Pol I inhibitor impaired the EMT/MET transitioning capability of tumor cells and significantly synergized with common chemotherapeutics. These observations also suggest that malignant cells might require certain ease to “ping pong” between epithelial and mesenchymal states, so that they could adapt themselves to the challenging microenvironment. Of note, some commonly used chemotherapeutics (Cisplatin, 5FU, Doxorubicin, etc), although primarily targeting DNA duplications, may also affect RiBi pathway by inhibiting rRNA processing(26). Therefore, the synergic effects with Pol I inhibitor varied among different combinations. Moreover, RiBi is a process dysregulated in most, if not all, cancers. Its involvement in EMT/MET process make it a feasible targeted pathway for treating patients with advanced breast cancer.

## ACKNOWLEDGEMENTS

We thank Dr. Jenny Xiang of the Genomics Resources Core Facility and Jason McCormick of the Flow Cytometry Core Facility for their professional advice. We thank Dr. Vivek Mittal and Dr. Nasser K. Altorki for their comments on this work. We thank Dr. Divya Ramchandani for technique support for experiments. This work was supported by National Cancer Institute (NCI) Grant NIH R01 CA205418 (D.G) and R01 CA244413 (D.G). This work was also supported by The Neuberger Berman Foundation Lung Cancer Research Center, a generous gift from Jay and Vicky Furman; and generous funds donated by patients in the Division of Thoracic Surgery to Dr. Altorki. The funding organizations played no role in experimental design, data analysis, or manuscript preparation.

## DATA ACCESS STATEMENT

Data generated in the study is either publicly available or by request to the corresponding authors.

## AUTHOR CONTRIBUTIONS

Y.B. and D.G. designed the experiments. Y.B., and Y.Z., performed the experiments. Y.B., D.G., J.S., Y.C., and S.W. performed bulk and single-cell RNA sequencing analyses. S.L and R.B. performed some animal works for experiments. D.G. supervised this study. Y.B. and D.G. wrote the manuscript. All authors discussed the results and conclusions drawn from the studies.

---

**Supplementary Fig S1.**
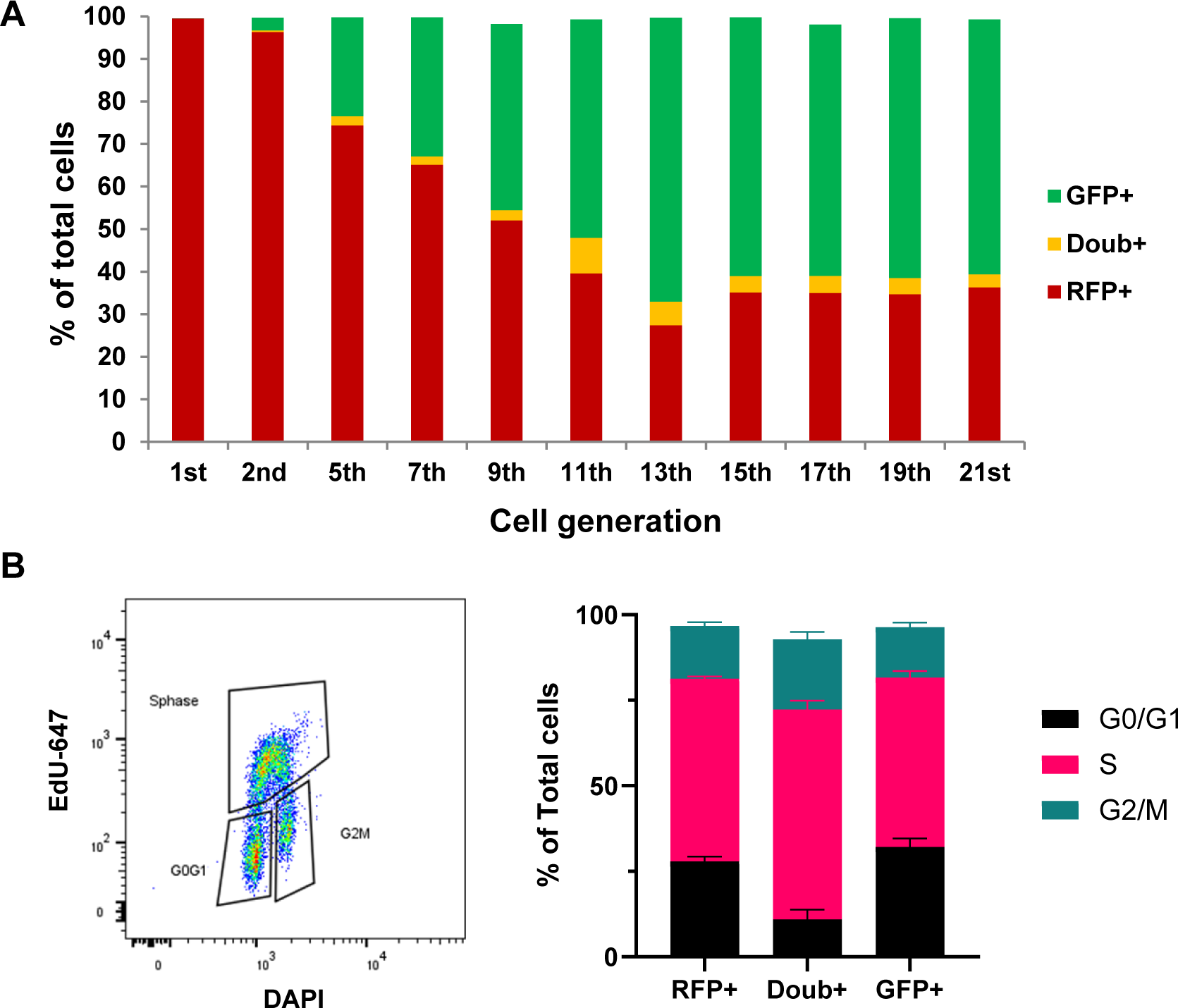
Characterization of fluorescent marker switch in Tri-PyMT cells. **A,** RFP+ Tri-PyMT cells were sorted via flow cytometry and cultured in DMEM with 10% FBS. Plot shows the percentage of RFP+, GFP+ and Doub+ cells at different generations of cell culture. The experiment was repeated twice with similar results. **B,** Cell cycle analysis of Tri-PyMT cells. Tri-PyMT cells (p7) composed by both RFP+ and GFP+ cells were cultured in growth medium. Cell cycle analysis was performed by using EdU incorporation and Click-chemistry. Cells in G0/G1, S and G2/M phases were quantified by flow cytometry. n =4, two-way Anova, P=0.0242, for S phase RFP+ vs GFP+; P < 0.0001, for S phases Doub+ vs RFP+ and Doub+ vs GFP+ cells.

**Supplementary Fig S2.**
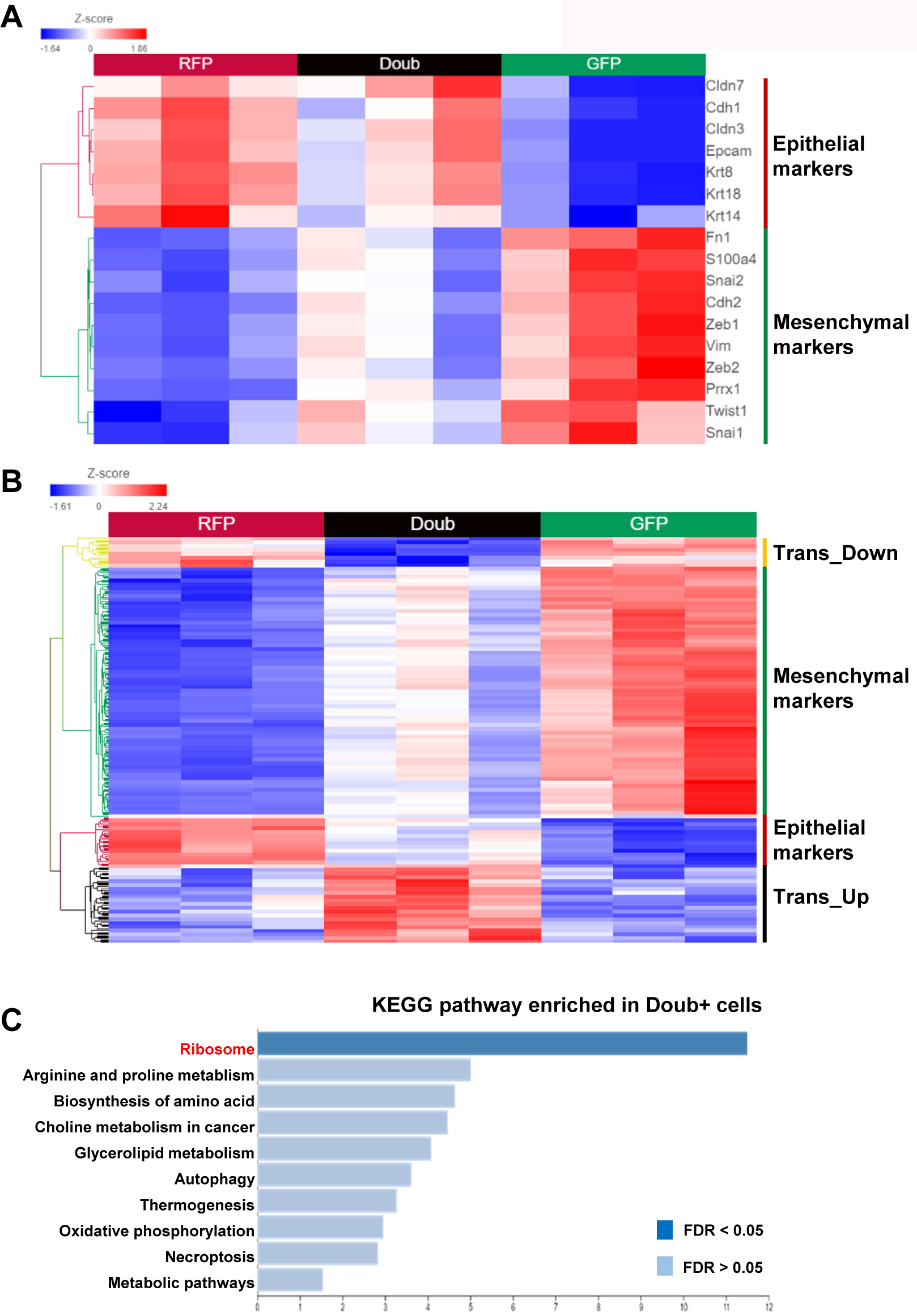
RNA-sequencing analysis of Tri-PyMT cells. RFP+ Doub+ and GFP+ Tri-PyMT cells were purified using flow cytometry sorting, total RNA was extracted and submitted for RNA-sequencing analysis. **A,** Heatmap of selected EMT marker genes in RFP+, Doub+ and GFP+ cells. Of note, the intermediate expression of both epithelial and mesenchymal markers in Doub+ cells. **B**, Heatmap of differentially expressed genes in Doub+ cells compared to RFP+ and GFP+ cells. In total, 213 genes are presented in 4 clusters, P<0.01, Doub+ vs RFP+ and GFP+. **C,** Gene set over-representative assay with upregulated genes in Doub+ cells showing the significant enrichment of Ribosome pathway (326 KEGG pathways analyzed in total).

**Supplementary Fig S3.**
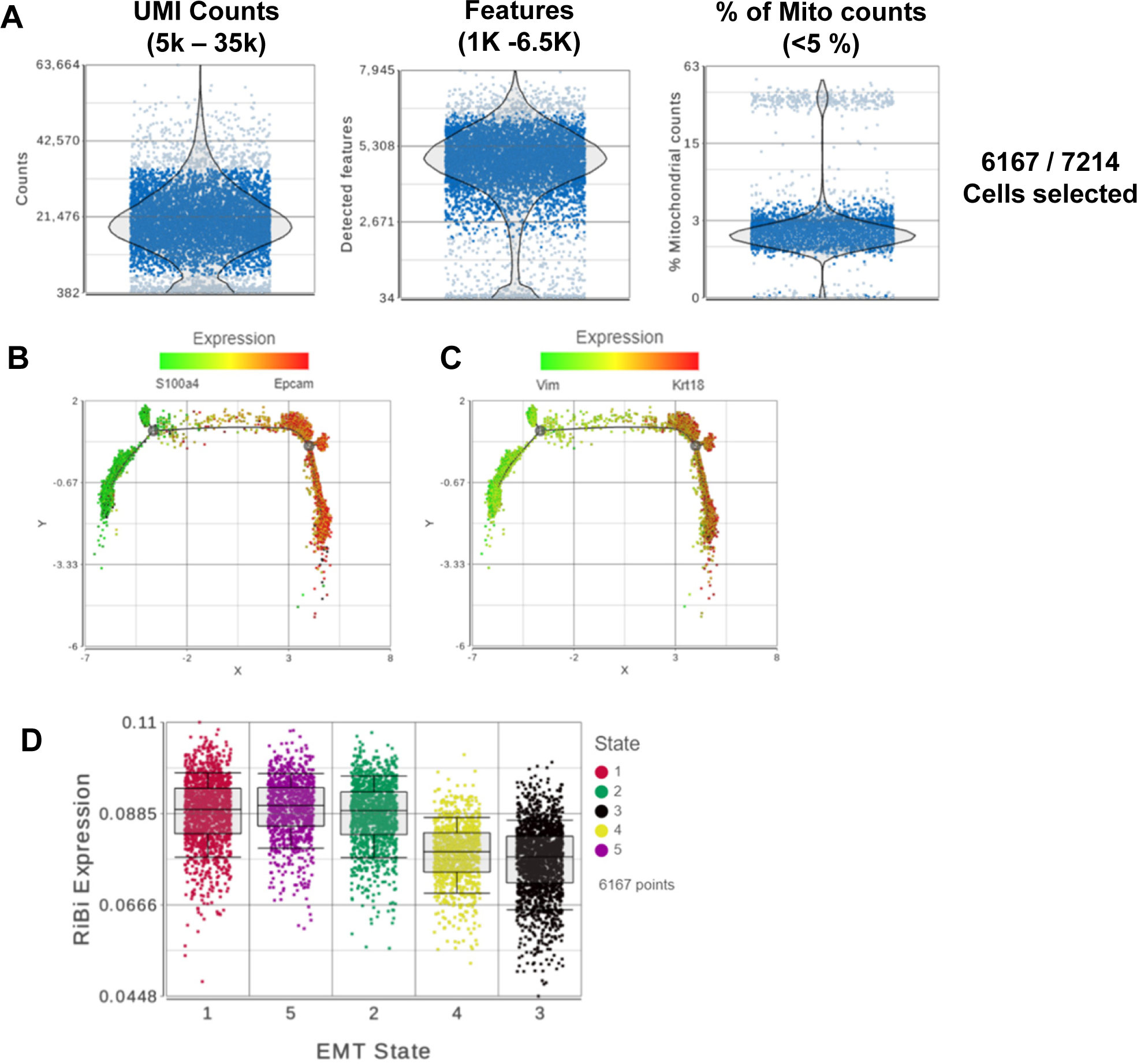
The scRNA-sequencing analysis of Tri-PyMT cells. RFP+, Doub+, and GFP+ Tri-PyMT cells were sorted via flow cytometry and submitted for sc-RNA sequencing analysis. **A.** Quality controls of the scRNA-seq. Plots show the single-cell QA/QC analyses including total counts of UMI, detected gene features, and the percentage of mitochondria genes. Totally, 6167 cells out of 7214 cells were filtered according to the criteria for downstream analyses. **B. C.** EMT trajectory analysis plots. Cell trajectory analysis was performed with filtered EMT-related genes using Monocle 2 model. Pairs of epithelial and mesenchymal marker genes, S100a4 vs Epcam (B) and Vim vs Krt18 (C), are used to highlight the EMT status of individual cells. **D.** Scatter plot shows the expression of Ribosome biogenesis genes (AUC values of RiBi pathway genes) according to EMT cell states. A trend of highest RiBi activation was detected in State 5 cells (p=0.072 for 5 vs 1, and p<0.01 for 5 vs 2, 3, 4, One-way ANOVA).

**Supplementary Figure S4:**
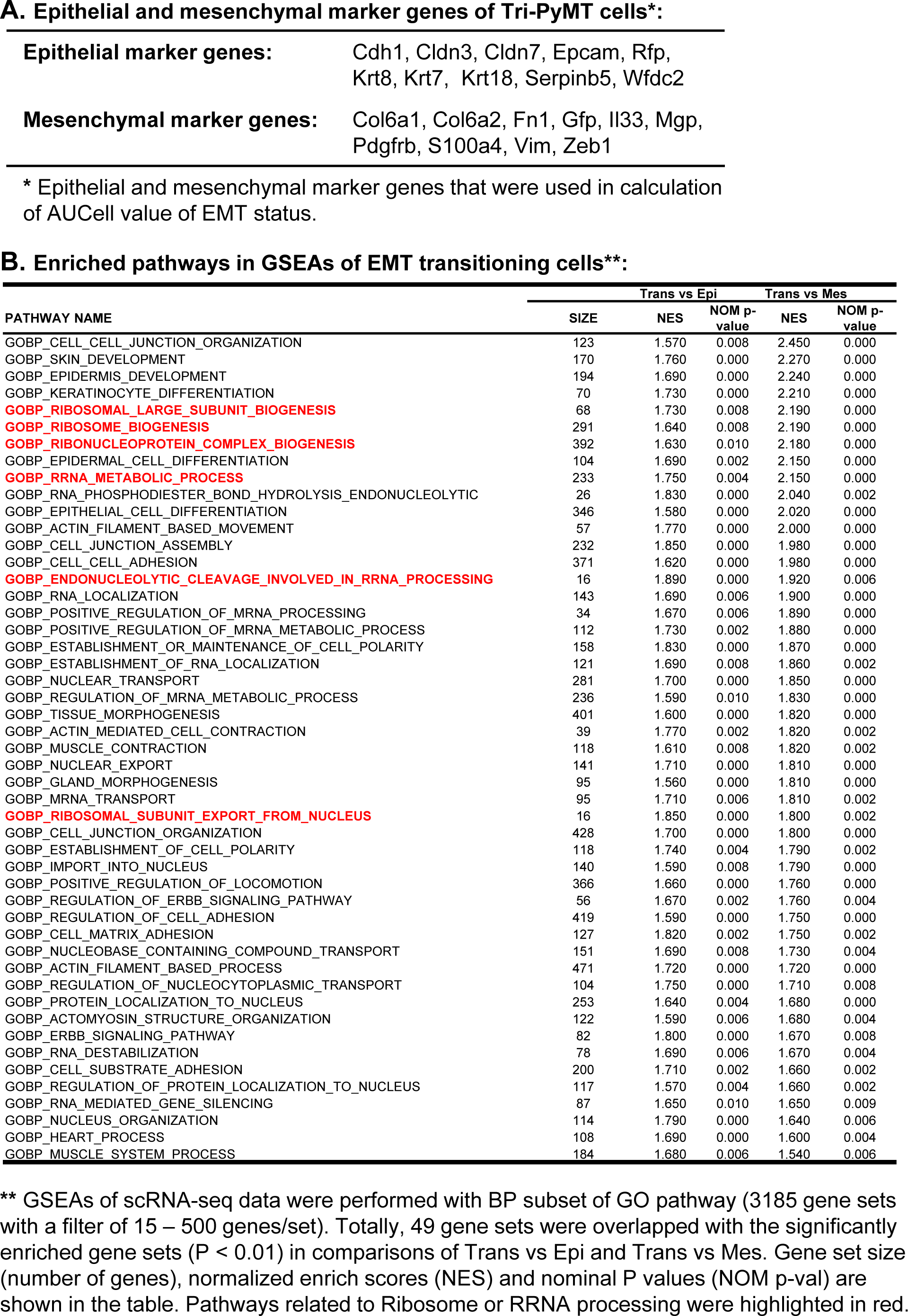

**Supplementary Fig S5.**
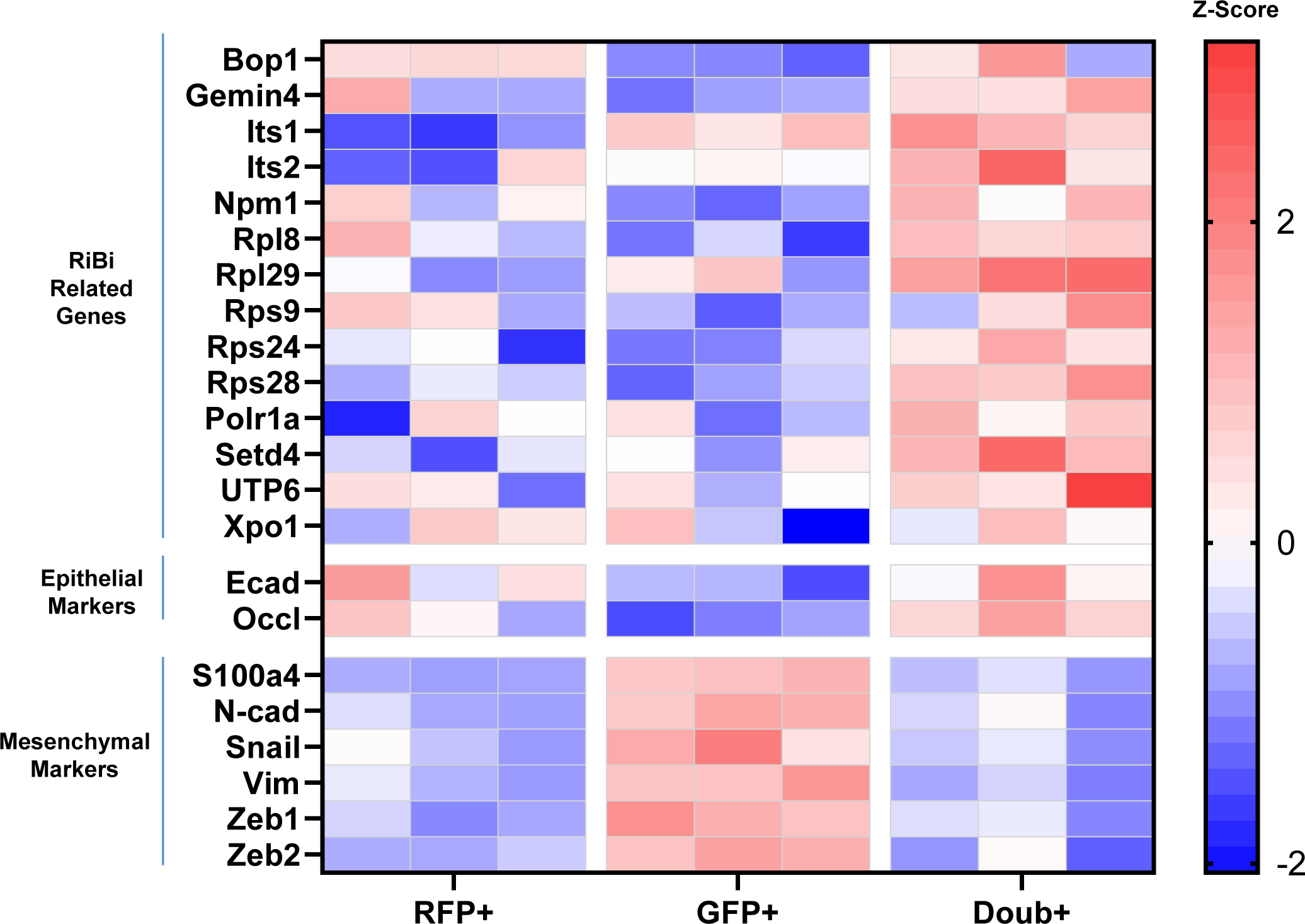
RT-PCR analysis of RiBi genes in Tri-PyMT cells. RFP+, Doub+ and GFP+ cells were purified from Tri-PyMT (p7) culture using flow cytometry sorting. Total RNA was extracted and analyzed by RT-PCR. The heatmap shows z-score of RiBi related genes, epithelial and mesenchymal marker genes expression with Gapdh as internal control, n=3.

**Supplementary Fig S6.**
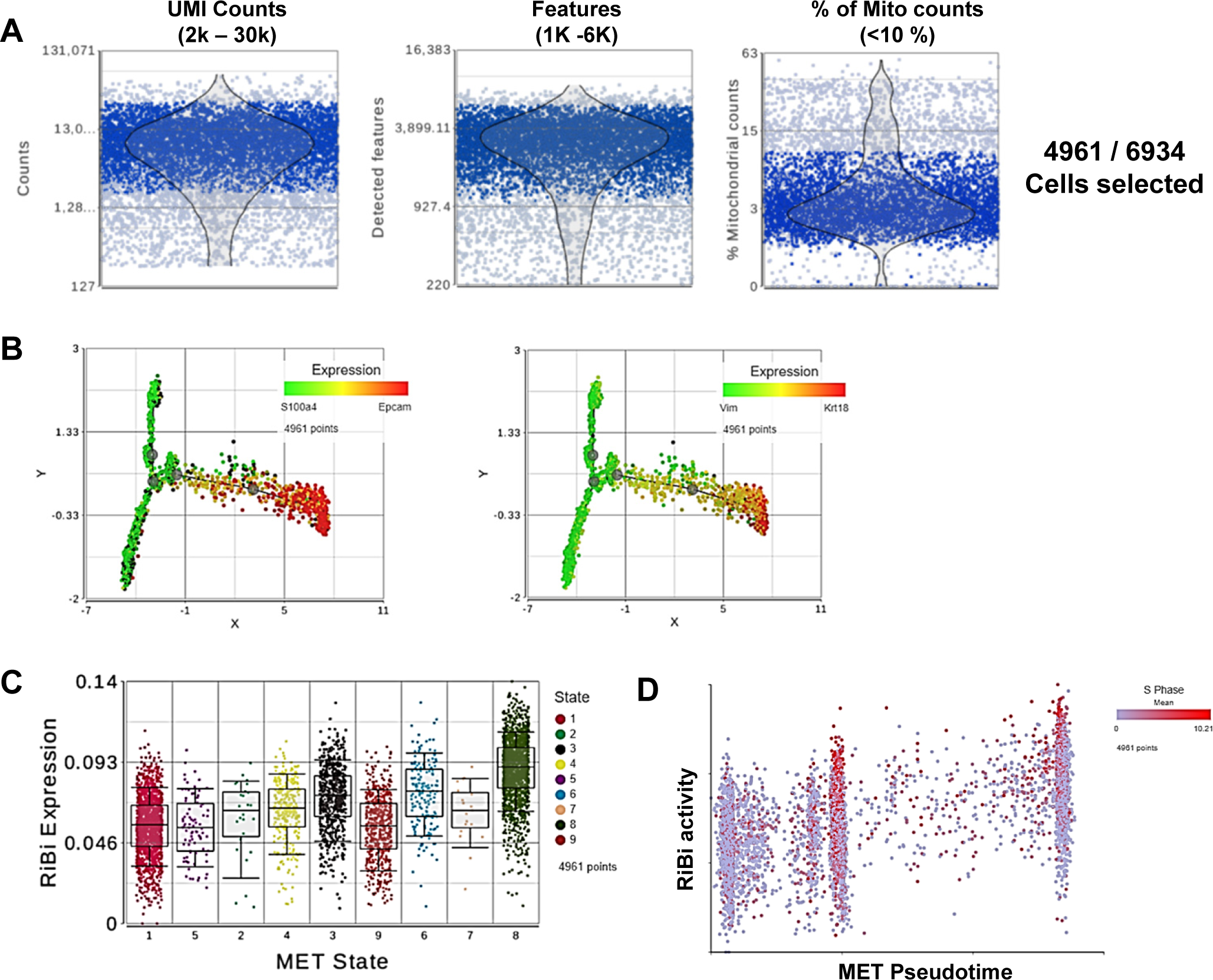
The scRNA-sequencing analysis of GFP+ Tri-PyMT cells. GFP+ Tri-PyMT cells, including Epcam+ and Epcam-cells were sorted from metastasis-bearing lungs and submitted for single cells RNA sequencing analysis. **A. Quality controls of the scRNA-seq.** Plots show the single-cell QA/QC analyses of the total counts of UMI, detected gene features and the percentage of mitochondria genes. Totally, 4961 cells out of 6934 cells were included in the downstream analyses. **B.** Plots of MET trajectory analysis. Cell trajectory analysis was performed with filtered EMTome genes using Monocle 2 model. Pairs of epithelial and mesenchymal marker genes, S100a4 vs Epcam (left) and Vim vs Krt18 (right) are used to highlight the MET status of individual cells. **C.** Elevated ribosome biogenesis pathway during MET. Cells were classified by their MET state. The activation of RiBi pathway was indicated by AUC value of genes in the ribosome biogenesis pathway. Of note, the highest RiBi expression in MET State 8 cells, which exhibit the most epithelial phenotypes. **D.** Scatter plot illustrates the correlation between RiBi activity and MET pseudotime. Proliferating cells (in S phase) are highlighted in red color.

**Supplementary Fig S7.**
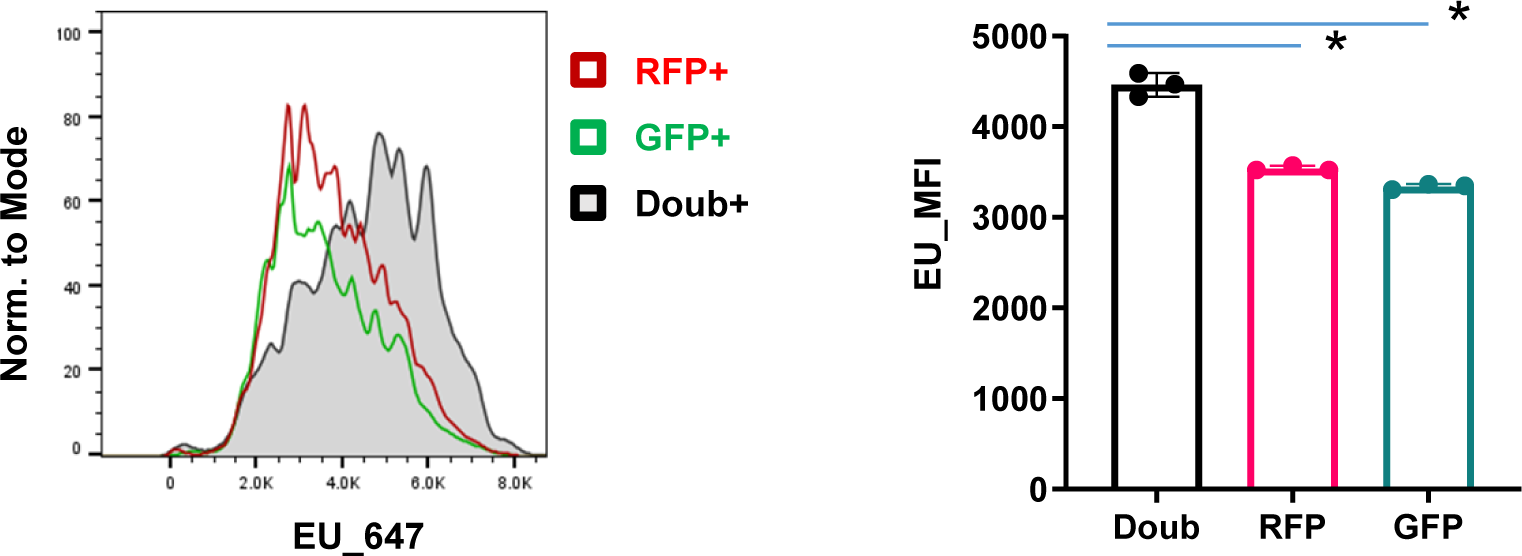
Elevated RNA transcription activity in the Doub+ Tri-PyMT cells. Tri-PyMT cells (p7) including both RFP+ and GFP+ cells were cultured in growth medium. EU incorporation assay was performed and detected with click chemistry using Alexa A647-azide. Labeled cells were analyzed by flow cytometry (Left). The median fluorescence intensity (MFI) of EU staining of RFP+, GFP+ and Doub+ cells were quantified, n =3, One-way ANOVA, *P < 0.0001, Doub+ vs RFP+; *P < 0.0001, Doub+ vs GFP+.

**Supplementary Fig. S8.**
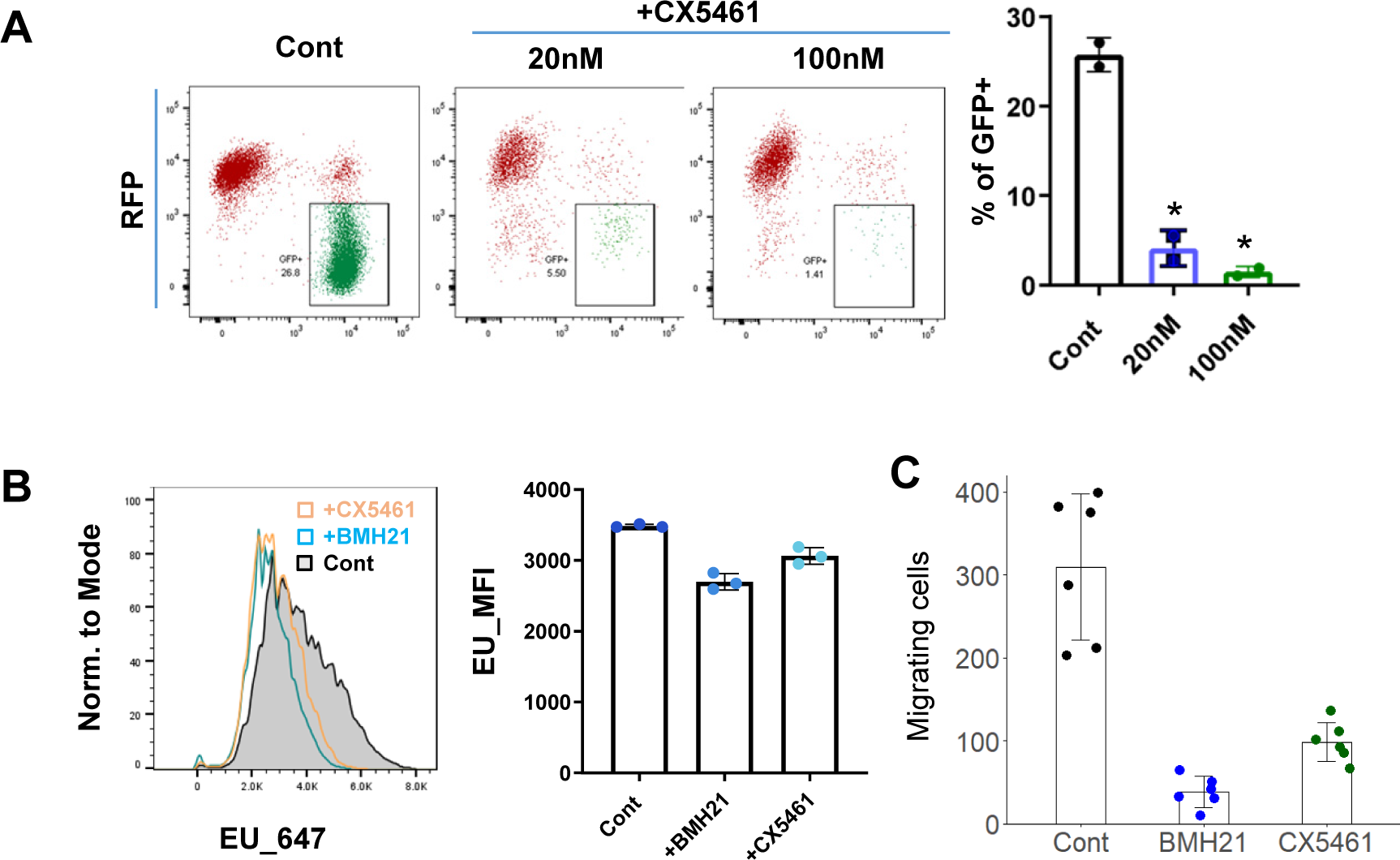
RiBi inhibition impairs RNA transcription activity and reduces EMT capability of tumor cells. **A,** Fluorescence switch in Tri-PyMT cells with CX5461 treatment. Flow cytometry plots exhibit the percentage of GFP+ Tri-PyMT cells after treatment with the Pol I inhibitor CX5461. Biological repeat, n=2, One-way ANOVA, *P=0.0015, 20nM vs Cont; *P=0.0011, 100nM vs Cont. **B,** EU incorporation assay. Tri-PyMT cells (p10) were treated with RiBi inhibitors, BMH21 at 100nM, or CX5461 at 20nM, for 2 days. EU incorporation assay was performed and detected with click chemistry using Alexa A647-azide. Labeled cells were analyzed by flow cytometry (Left). The median fluorescence intensity (MFI) of EU staining of cells were quantified, n =3, One-way ANOVA, P = 0.0001, +BMH21 vs Cont; P = 0.0032, +CX5461 vs Cont. **C,** Cell migration assay. Tri-PyMT cells (p7) were seeded in the inserts of migration plate and treated BMH21 (100nM) or CX5461 (20nM) for 5 days. Cell migration was induced by FBS gradient for 24 hours. Migrating cells were counted from the bottom of the insert. n=6, One-way ANOVA, * P < 0.001 BMH21 vs Cont and CX5461 vs Cont.

**Supplementary Fig S9.**
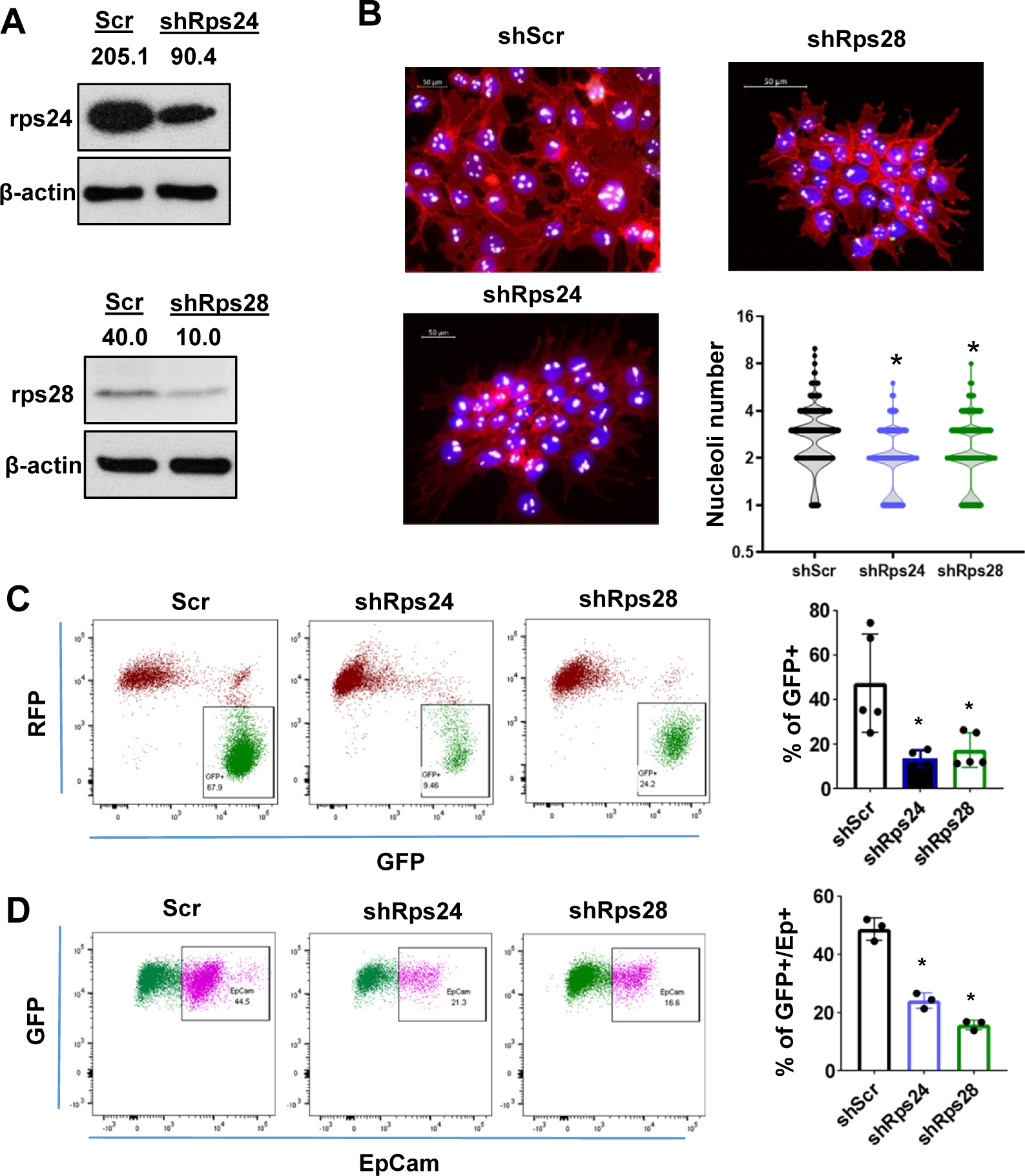
RiBi modulating by knocking down ribosome proteins reduces EMT/MET capability of tumor cells. **A,** Western blots show the knockdown expression of Rps24 and Rps28 in targeted Tri-PyMT cells. The numbers indicate relative intensity of the band normalizing to the β-actin band of the sample. **B,** Impact on nucleoli numbers by Rps knockdown in RFP+ Tri-PyMT cells. Fluorescent images show the reduced number of nucleoli in Rps24 and Rps28 knockdown cells. n = 623 cells (shScr), 490 cells (shRps24), 553 cells (shRps28). One-way ANOVA with Dunnett multiple comparisons, *P < 0.0001 for shScr vs shRps24 and shScr vs shRps28. **C,** Flow cytometry plots show the percentage of GFP+ cells in Rps24 and Rps28 knockdown Tri-PyMT cells. 4 biological repeats, One-way ANOVA, *P=0.0085, shScr vs. shRps24, *P=0.0117, shScr vs. shRps28. **D,** Flow cytometry plots show the Epcam+/GFP+ cells in Rps24 or Rps28 knockdown Tri-PyMT cells. 3 biological repeats, one-way ANOVA, *P=0.0055, shScr vs. shRps24, *P=0.0124, shScr vs. shRps28.

**Supplementary Fig S10.**
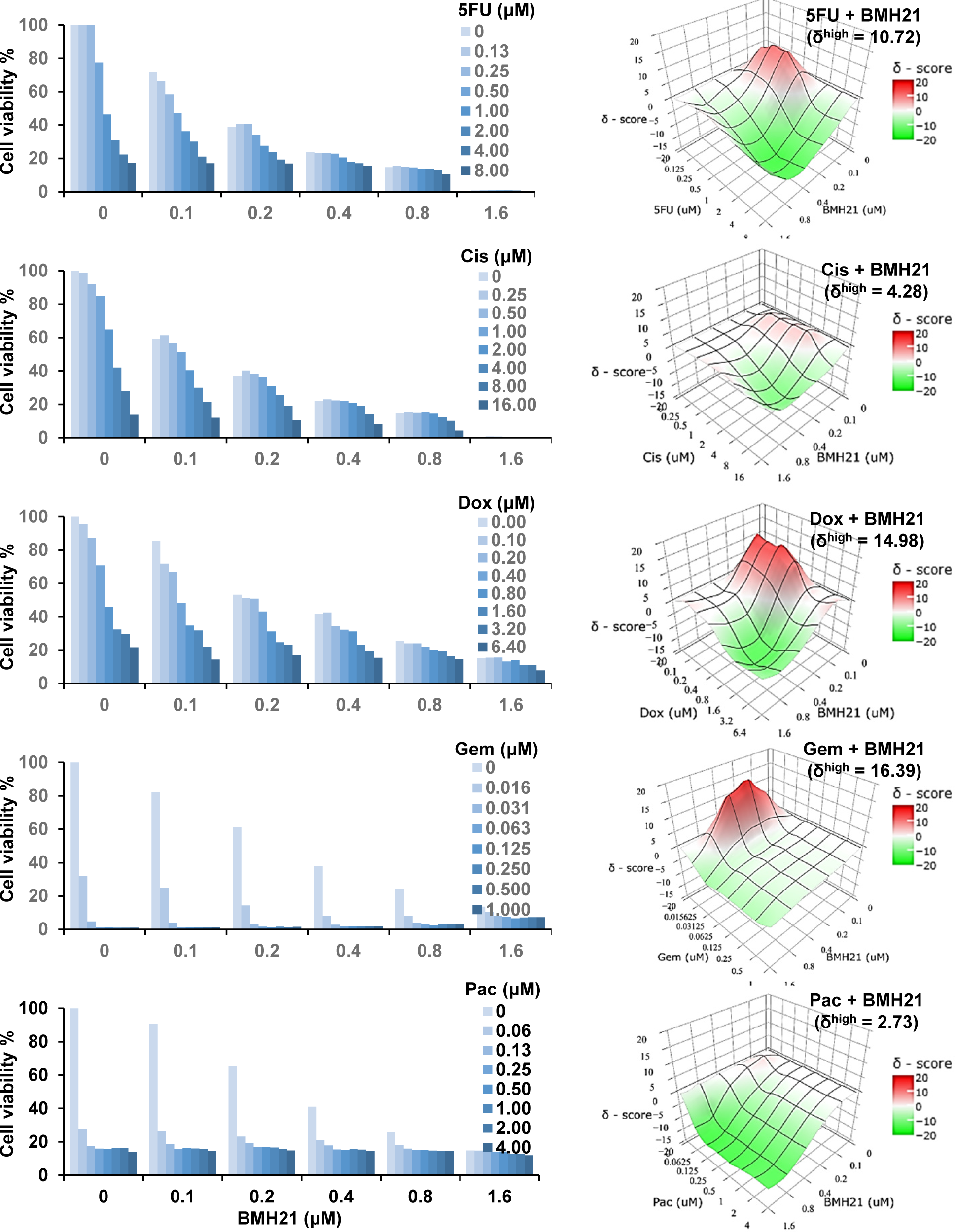
Combination therapy of RNA Pol I inhibitor and chemo drugs. **Left**, Bar graphs display Tri-PyMT cell viability following combination treatment with BMH21 and chemotherapeutic drugs at various concentrations. Bars represent the mean value of duplicated treatments, n=2 wells/treatment. Right, Synergy plots of BMH21 and chemotherapeutic drugs. Synergistic scores (δ) for each combination were calculated using the ZIP model in SynergyFinder 3.0. The highest δ numbers among combinations is shown.

**Supplementary Fig S11.**
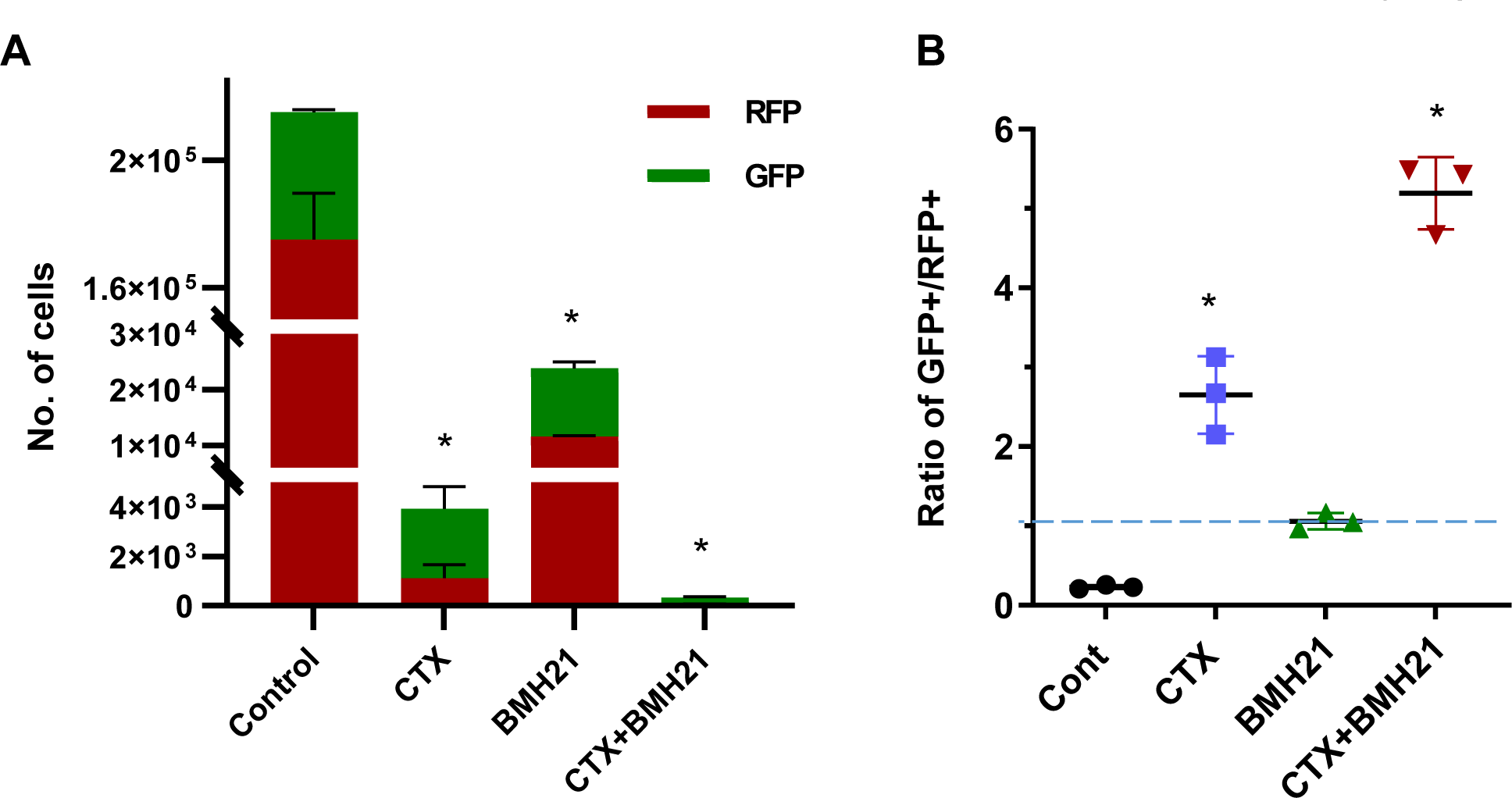
Differential impact of Pol I inhibitor and chemo drug on epithelial and mesenchymal Tri-PyMT cells *in vivo*. The same number of RFP+ and GFP+ Tri-PyMT cells were injected in animals via tail vein. Animals were treated CTX (100mg/kg, once a week, i.p.), BMH21 (20mg/kg, 5 times a week, i.p.) or both in combination. Number of RFP+ and GFP+ cells were quantified by flow cytometry at day 21 after inoculation. **A,** Plots show the number of RFP+ and GFP+ Tri-PyMT cells in the lung treated with CTX and/or BMH21. n=3 mice/treatment, One way ANOVA, *P<0.0001 for Cont vs CTX, Cont vs BMH21 and Cont vs CTX+BMH21. **B,** The ratio of GFP+ vs RFP+ cells in the lung treated with CTX and/or BMH21. n=3 mice/treatment, One way ANOVA, *P=0.0001 for Cont vs CTX; *P=0.0661 for Cont vs BMH21 and *P<0.0001 for Cont vs CTX+BMH21.

**Supplementary Fig S12.**
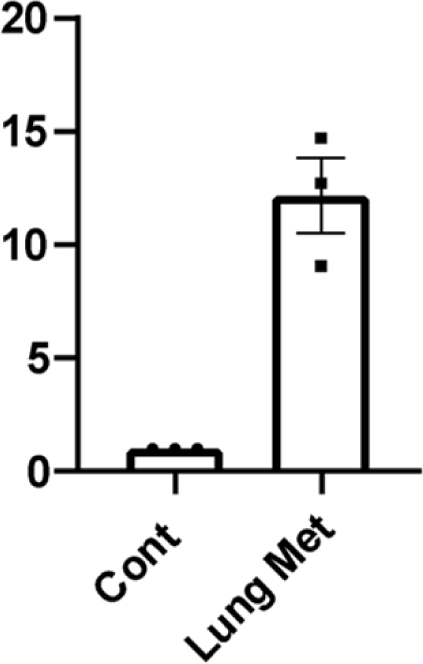
E-cadherin expression by LM2 cells in lung metastases. qRT-PCR analysis of Ecadherin expression in LM2 cells isolated from lung metastases. LM2 cells in culture served as control. GAPDH served as internal control for PCR. n=3 experiments, unpaired t test, * P=0.0025.

**Supplementary Fig 13.**
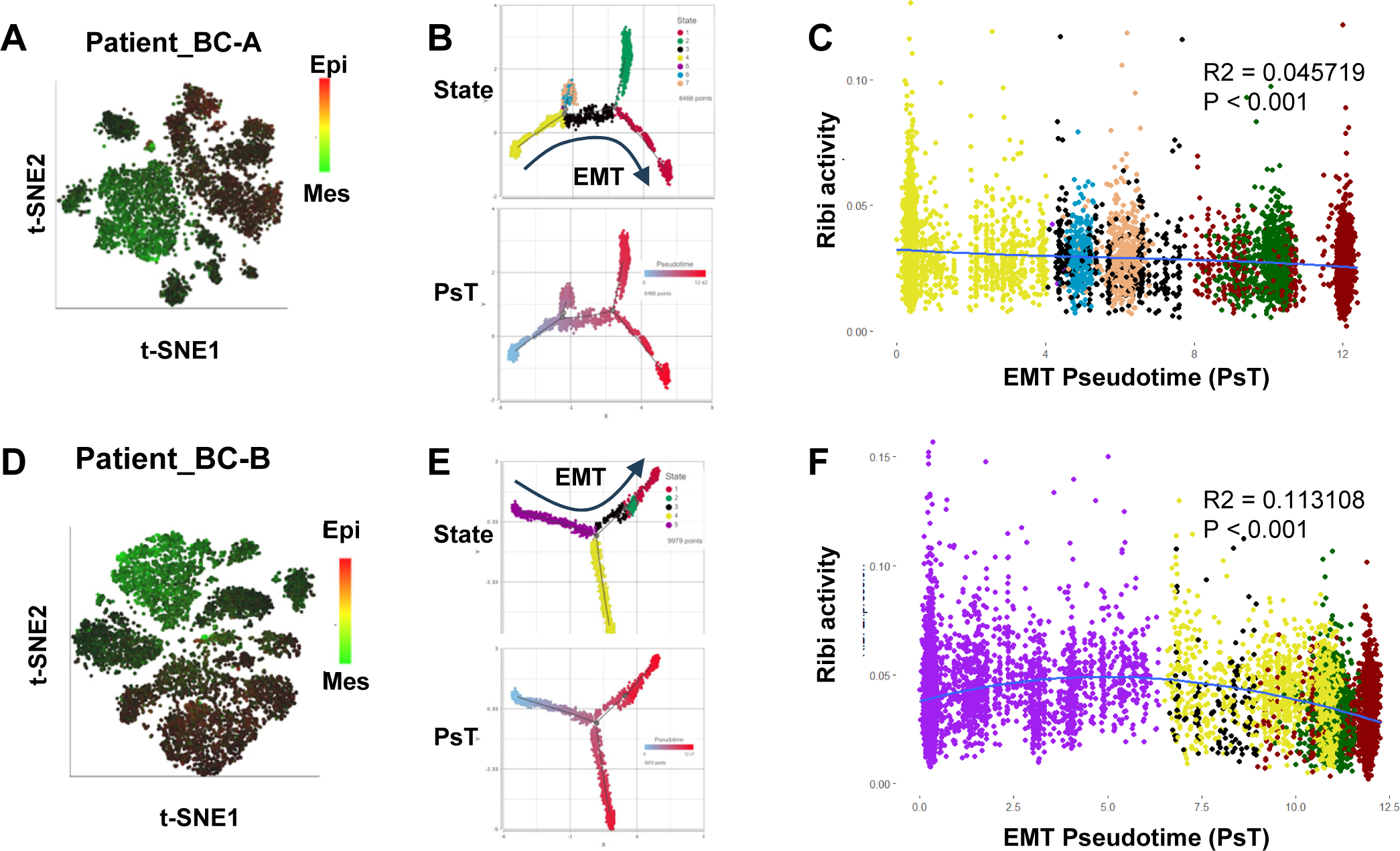
Correlation of ribosome biogenesis pathway and EMT status in human breast cancer cells. Single-nucleus cell RNA sequencing analysis of primary tumor cells from breast cancer patients (BC-A and BC-B, GEO database, GSE 198745). **A,D,** The *t*-SNE plots show the heterogeneity in EMT statuses of tumor cells with AUC values of epithelial genes (in red) and mesenchymal genes (in green). **B,E,** Cell trajectory analysis was performed with filtered EMT-related genes (EMTome genes) using Monocle 2 model. Cell states were identified with the most epithelial cells as root for calculation of EMT pseudotime (PsT). **C,F,** The scatter plot displays the correlation of Ribi activity to EMT pseudotime. The polynomial regression line (order = 3) highlights the higher RiBi activity in cells toward epithelial phenotypes.

**Supplementary Fig S14.**
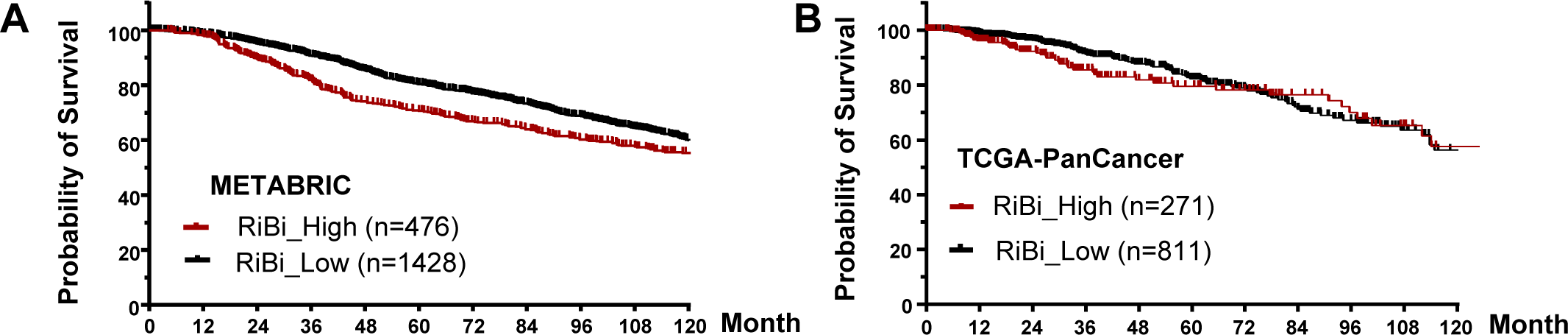
Survival curves of breast cancer patients with different RiBi activities. Gene expression data of METABRIC and TCGA-PanCancer was obtained from CBioPortal. Patients were grouped according to the average z-score of RiBi gene expression, RiBi High (>1) and RiBi Low (RiBi score <0.5). Upper, METABRIC dataset, Gehan-Breslow-Wilcoxon test *P*-value = 0.0019. Lower, TCGA data, Gehan-Breslow-Wilcoxon test *P*-value = 0.0225.

**Supplementary Fig S15.**
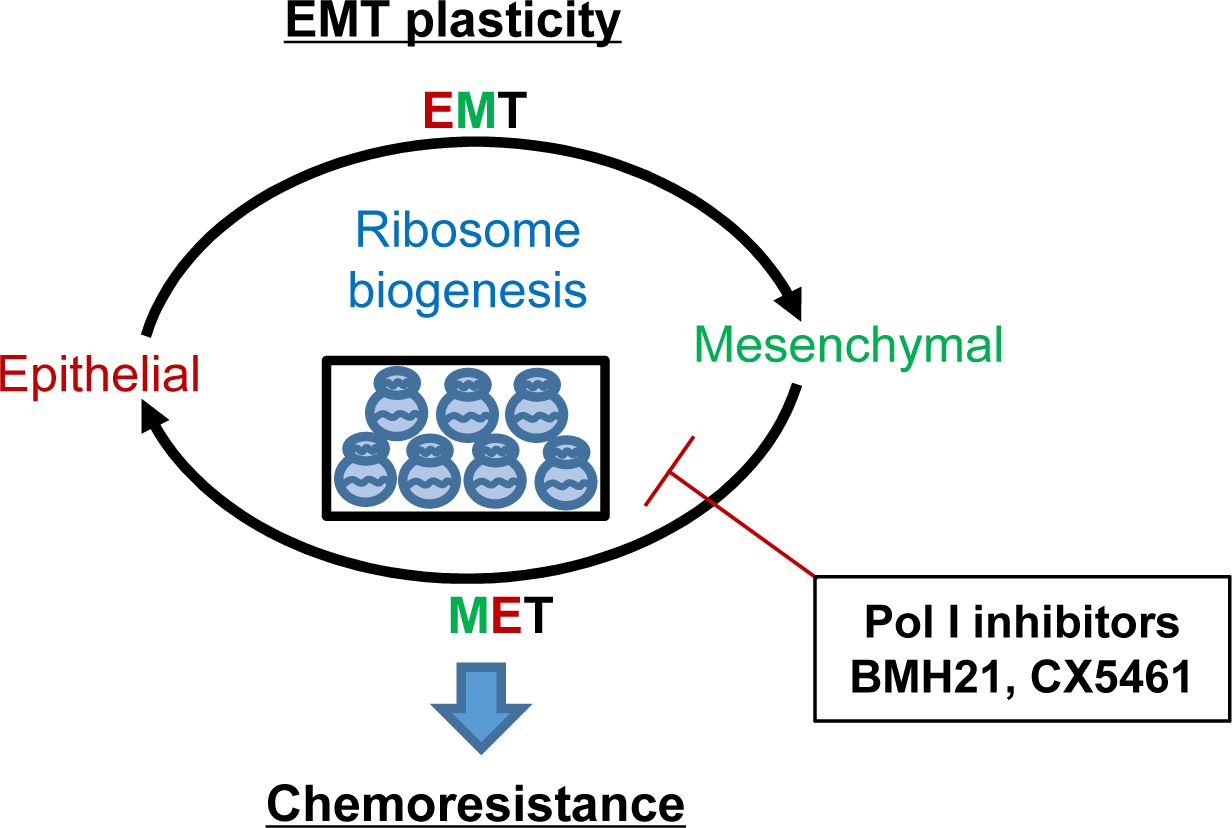
Working model of ribosome biogenesis in EMP plasticity. Tumor cells exhibit plasticity by undergoing EMT (epithelial-mesenchymal transition) and MET (mesenchymal-epithelial transition), contributing to the development of resistance to chemotherapies. Elevated ribosome biogenesis is essential for maintaining this EMT plasticity. Inhibition of ribosome biogenesis by RNA Pol I inhibitors synergizes with chemotherapeutic drugs to overcome resistance development.

